# Antimicrobial resistance prevalence, rates of hospitalization with septicemia and rates of mortality with sepsis in adults in different US states

**DOI:** 10.1101/404137

**Authors:** Edward Goldstein, Derek R. MacFadden, Zeynal Karaca, Claudia A. Steiner, Cecile Viboud, Marc Lipsitch

## Abstract

**Objectives:** Rates of hospitalization with sepsis/septicemia and associated mortality in the US have risen significantly during the last two decades. Antibiotic resistance may contribute to the rates of sepsis-related outcomes through lack of clearance of bacterial infections following antibiotic treatment during different stages of infection. However, there is limited information about the relation between prevalence of resistance to various antibiotics in different bacteria and rates of sepsis-related outcomes.

**Methods:** For different age groups of adults (18-49y,50-64y,65-74y,75-84y,85+y) and combinations of antibiotics/bacteria, we evaluated associations between state-specific prevalence (percentage) of resistant samples for a given combination of antibiotics/bacteria among catheter-associated urinary tract infections in the CDC Antibiotic Resistance Patient Safety Atlas data between 2011-2014 and rates of hospitalization with septicemia (ICD-9 codes 038.xx present on the discharge diagnosis) reported to the Healthcare Cost and Utilization Project (HCUP), as well as rates of mortality with sepsis (ICD-10 codes A40-41.xx present on death certificate).

**Results:** Among the different combinations of antibiotics/bacteria, prevalence of resistance to fluoroquinolones in *E. coli* had the strongest association with septicemia hospitalization rates for individuals aged over 50y, and with sepsis mortality rates for individuals aged 18-84y. A number of positive correlations between prevalence of resistance for different combinations of antibiotics/bacteria and septicemia hospitalization/sepsis mortality rates in adults were also found.

**Conclusions:** Our findings, as well as our related work on the relation between antibiotic use and sepsis rates support the association between resistance to/use of certain antibiotics and rates of sepsis-related outcomes, suggesting the potential utility of antibiotic replacement.

## 1. Introduction

Rates of hospitalization with septicemia (ICD-9 codes 038.xx) as well as associated mortality and monetary costs have been rising rapidly during the past decades in the US [1-4]. While changes in diagnostic practices have contributed to the rise in the rates of hospitalization with septicemia/sepsis in the diagnosis [5,6], those changes cannot fully explain that rise in hospitalization rates, particularly prior to 2010 [7]. Indeed, trends in the rates of US hospitalizations with any diagnosis of sepsis between 2003-2009 closely resemble the trends in the rates of hospitalizations that involved infection and the use of mechanical ventilation (Figure 1 in [7]). Additionally, rates of hospitalization with severe sepsis in the US were growing quite rapidly between 2008-2012, and the percent of those hospitalizations with multiple organ failure was also increasing [8], supporting the notion that the growth in the volume of those hospitalizations was genuine.

Antibiotic resistance contributes to the rates of septicemia/sepsis hospitalization and associated mortality. The relation between infections with antibiotic-resistant bacteria and survival for sepsis, including the effect of initially appropriate antibiotic therapy (IAAT) is suggested by a number of studies [9]. However, less is known about the relation between prevalence of antibiotic resistance and rates of hospitalization with septicemia. Antimicrobial resistance and use can contribute to the volume of hospitalizations associated with bacterial infections, including septicemia and severe sepsis, through several mechanisms. Importantly, antibiotic resistance can facilitate the progression to a severe disease state when infections not cleared by antibiotics prescribed during both the outpatient and the inpatient treatment eventually devolve into sepsis and associated lethal outcomes. Some of the more direct evidence for the relation between antibiotic resistance and subsequent hospitalization with severe infections, including bacteremia/sepsis is described in [10,11]. For example, prevalence of resistance to certain antimicrobials, including ciprofloxacin and trimethoprim, was found to be significantly higher in bacteremia outcomes associated with urinary tract infections (UTIs) than in the population of patients with UTIs in England [10], suggesting treatment failure to be an important factor behind the volume of UTI-associated bacteremia. Another study documented an association between trimethoprim resistance, trimethoprim use in UTI treatment and subsequent UTI-related bacteremia in England [12], whereas association between amoxicillin use in England and prevalence of trimethoprim resistance (likely modulated by the high rates of cross-resistance) was shown in [13]. The latter represents an example of how resistance to/use of one antibiotic affects the prevalence of other resistance phenotypes, or pathogenic organisms, with additional examples being the relation between fluoroquinolone use/resistance and (i) methicillin-resistant staphylococcus aureus (MRSA) infections [14,15]; (ii) extended-spectrum beta-lactamese (ESBL) producing infections [16,17]; (iii) *Clostridium difficile* (*C. difficile*) infections [18].

In our related work [19,20], we have studied the relationship between population-level prescribing of different antibiotic types/classes and rates of septicemia/bacteremia hospitalization and sepsis mortality, showing an association between the use of penicillins and (a) rates of septicemia hospitalizations and sepsis mortality in older US adults; (b) rates of *Escherichia coli* (*E. coli*) and methicillin-sensitive staphylococcus aureus (MSSA) bacteremia in England. This work represents another population level study, examining the relationship between prevalence of resistance for different combinations of bacteria/antibiotics and rates of hospitalization with septicemia and mortality with sepsis in US adults. Specifically, we examine associations between state-specific prevalence of antibiotic resistance for different combinations of antibiotics/bacteria in hospitalized elderly and non-elderly adults documented in the US CDC National Healthcare Safety Network (NHSN) Antibiotic Resistance Patient Safety Atlas [21], and (a) state-specific rates of hospitalization with septicemia (ICD-9 codes 038.xx present on a discharge diagnosis) in different age groups of adults recorded in the Healthcare Cost and Utilization Project (HCUP) data [22]; (b) state-specific rates of mortality with sepsis (ICD-10 codes A40-41.xx present on a death certificate) in different age groups of adults recorded in the US CDC Wonder data [23]. The aim of this work, as well as of the analyses in [19,20] is to provide further evidence that resistance to/use of certain antibiotics is associated with sepsis-related outcomes.

## 2. Materials and Methods

### 2.1 Hospitalizations with septicemia and mortality with sepsis

We used weekly data between 2011-2012 on counts of hospitalizations with a septicemia diagnosis (both primary and contributing, ICD-9 codes 038.xx) from the State Inpatient Databases of the Healthcare Cost and Utilization Project (HCUP), maintained by the Agency for Healthcare Research and Quality (AHRQ) through an active collaboration [22]. We will henceforth call these hospitalizations septicemia hospitalizations, even though some of them may involve low levels of bloodstream bacterial infection. This database contains hospital discharges from community hospitals in participating states. Forty-two states reported septicemia hospitalization data between 2011-2012 for each of the five adult age subgroups included in our analyses: (18-49y, 50-64y, 65-74y, 75-84y, 85+y). Those states are Alaska, Arkansas, Arizona, California, Colorado, Connecticut, Florida, Georgia, Hawaii, Iowa, Illinois, Indiana, Kansas, Kentucky, Louisiana, Massachusetts, Maryland, Minnesota, Missouri, Montana, North Carolina, North Dakota, Nebraska, New Jersey, New Mexico, Nevada, New York, Ohio, Oklahoma, Oregon, Rhode Island, South Carolina, South Dakota, Tennessee, Texas, Utah, Virginia, Vermont, Washington, Wisconsin, West Virginia, Wyoming.

Data on annual state-specific mortality with sepsis (ICD-10 codes A40-A41.xx representing either the underlying or a contributing cause of death) for the 50 US states and District of Columbia between 2013-2014 for different age groups of adults (18-49y, 50-64y, 65-74y, 75-84y, 85+y) were extracted from the US CDC Wonder database [23]. *Average annual* septicemia hospitalization rates between 2011-2012, as well as the average annual sepsis mortality rates between 2013-2014 per 100,000 individuals in each age group/state were then calculated from the septicemia hospitalization/sepsis mortality data and population estimates, taken to be the the annual, July 1 population estimates in [24]. Finally, we note that the ICD-9 code 038.xx (septicemia) has no exact analogue in the ICD-10 coding system. Nonetheless, the vast majority of hospitalizations with septicemia have the 038.9 code (unspecified septicemia) on the discharge diagnosis, with that code translating to A41.9 (sepsis, unspecified organism) in the ICD-10 coding system, with the latter code appearing in the vast majority of deaths with sepsis (ICD-10 codes A40-41.xx listed on the death certificate).

*Ethics statement:* Aggregate hospitalization and mortality data were used in our analyses, with no informed consent from participants sought.

### 2.2 Antibiotic resistance prevalence

We extracted data on the prevalence of antibiotic resistance for bacterial specimens collected from hospitalized patients in the US between 2011-2014 from the US CDC National Healthcare Safety Network (NHSN) Antibiotic Resistance Patient Safety Atlas (AR Atlas) [21]. Those data are stratified by age group (<1y,1-18y,19-64y,65+y), state, year, infection type (CAUTI/CLABSI/SSI, [21]), and combination of bacteria/antibiotics (31 combinations documented in [21]). Resistance prevalence in [21] varies by type of infection, making the combination of different types of infection into the analysis problematic. Moreover, for a given combination of bacteria/antibiotics, and a given type of infection, only a fraction of states reported the corresponding resistance pattern [21], with that fraction generally being highest for catheter-associated urinary tract infections (CAUTIs). For each combination of bacteria/antibiotics, age group of adults (19-64y or 65+y), and state that reported data on CAUTI samples between 2011-2014 for the given age group/combination of bacteria/antibiotics, the corresponding state-specific *prevalence of resistance* was defined as the percent of tested CAUTI samples collected between 2011-2014 for the given age group/state containing the corresponding bacteria that were resistant (or have tested as either intermediate or resistant – see [21]) for the corresponding antibiotics. The four-year aggregation was done due to low (or non-specified) yearly counts in a number of states.

### 2.3 Correlation analyses

For each age group of adults: (18-49y, 50-64y, 65-74y, 75-84y, 85+y), and a combination of bacteria/antibiotics, we have examined correlations, both linear (Pearson) and Spearman (Supporting Information), between (i) the state-specific average annual septicemia hospitalization rates per 100,000 individuals in the given age group of adults, 2011-2012; (ii) the state-specific average annual sepsis mortality rates per 100,000 individuals in the given age group of adults, 2013-2014 and the state-specific prevalence of resistance in CAUTI samples (see the previous subsection), 2011-2014 for the given combination of bacteria/antibiotics among the non-elderly or elderly adults correspondingly. For each age group and sepsis-related outcome (septicemia hospitalizations or sepsis mortality), the above correlations are computed for those combinations of antibiotics/bacteria for which at least 10 states reported the corresponding data. We note that no septicemia hospitalization data beyond 2012 were available for this study, and that we used the two most recent years (2013-2014) for the mortality data due to potential changes in coding for sepsis mortality on death certificates [25]. We also note that CAUTIs represent only a small fraction of all septicemia hospitalizations/subsequent sepsis mortality. Nonetheless, we use prevalence of resistance in the CAUTI samples as a proxy for the statewide prevalence of resistance in different settings, under the premise that this source of noise should generally bias the correlation estimates towards the null, rather than create spurious associations.

## 3. Results

Figures 1-5 show the linear (Pearson) correlations between the state-specific prevalence (percentages) of antibiotic resistance for the different combinations of antibiotics/bacteria in the age-specific CAUTI samples in the CDC AR Atlas data [21] between 2011-14 and the state-specific average annual rates of hospitalizations between 2011-12 with septicemia in either the principal or secondary discharge diagnosis recorded in the HCUP data [22] per 100,000 individuals in the corresponding age group. Figures 6-10 present the linear correlations between the state-specific prevalence of antibiotic resistance [21] and rates of sepsis mortality [23] between 2013-14 in different age groups of adults. All the correlations are presented for those combinations of antibiotics/bacteria and age group for which at least 10 states reported the corresponding data. More detailed results of the correlation analyses, including Spearman correlations between prevalence of resistance and rates of sepsis-related outcomes are presented in the Supporting Information.

**Figure 1:**
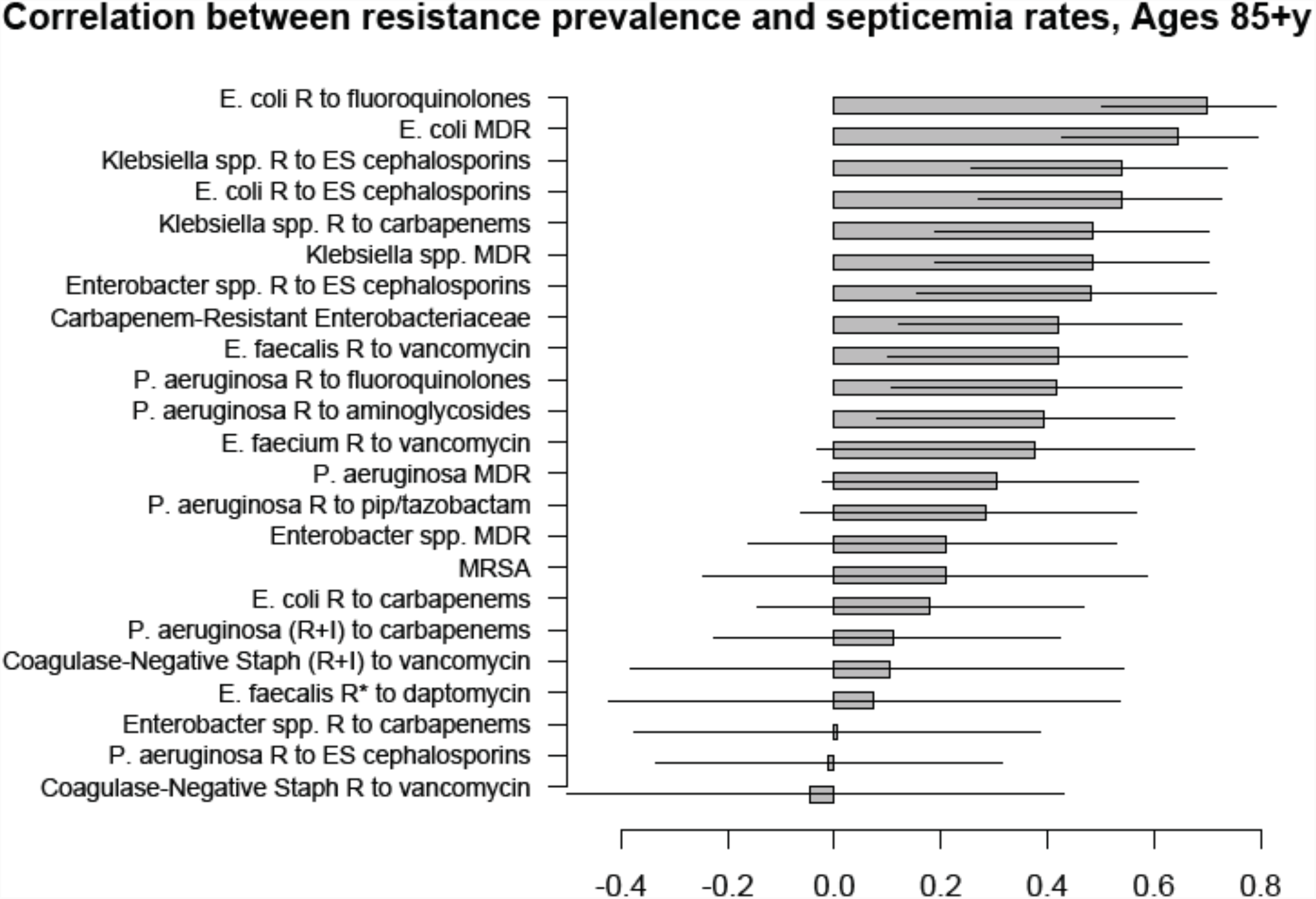
Correlation between state-specific prevalence (percentages) of resistance for different combinations of antibiotics/bacteria in CAUTI samples from hospitalized adults aged 65+y in the CDC AR Atlas data [21] between 2011-14 and state-specific average annual rates per 100,000 individuals aged 85+y of septicemia hospitalizations (principal or secondary diagnosis) recorded in the HCUP data [22].

**Figure 2:**
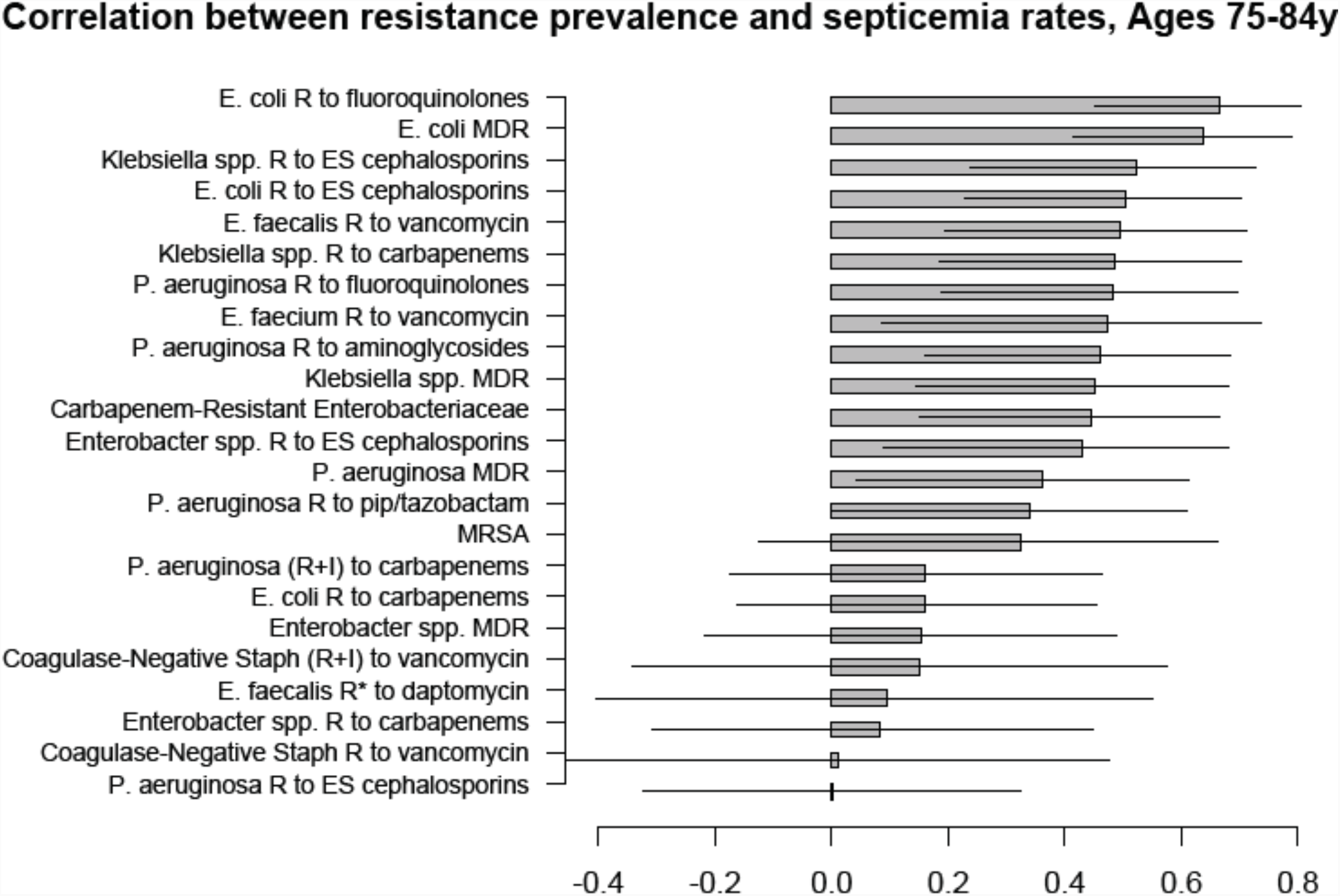
Correlation between state-specific prevalence (percentages) of resistance for different combinations of antibiotics/bacteria in CAUTI samples from hospitalized adults aged 65+y in the CDC AR Atlas data [21] between 2011-14 and state-specific average annual rates per 100,000 individuals aged 75-84y of septicemia hospitalizations (principal or secondary diagnosis) recorded in the HCUP data [22].

**Figure 3:**
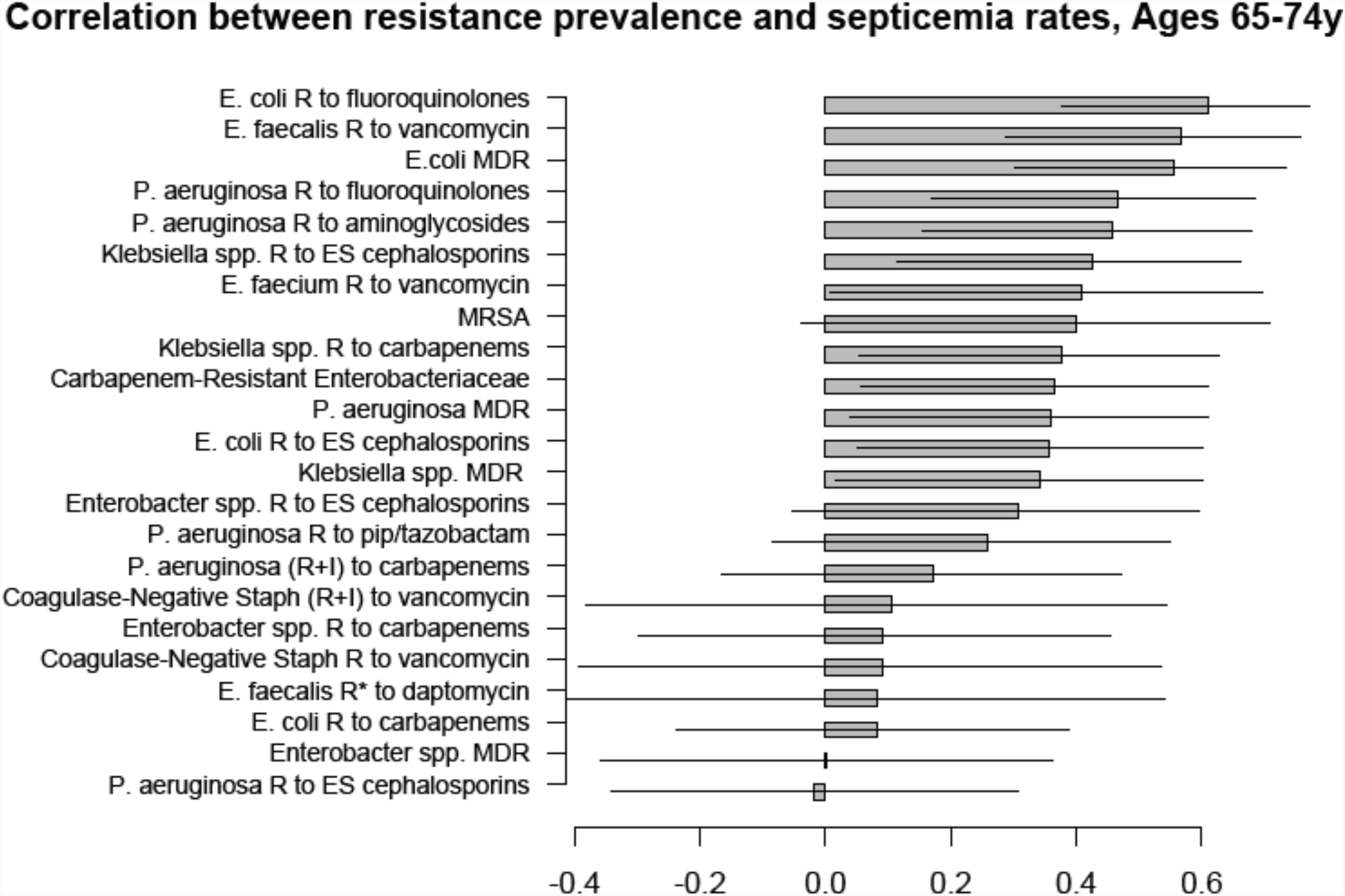
Correlation between state-specific prevalence (percentages) of resistance for different combinations of antibiotics/bacteria in CAUTI samples from hospitalized adults aged 65+y in the CDC AR Atlas data [21] between 2011-14 and state-specific average annual rates per 100,000 individuals aged 65-74y of septicemia hospitalizations (principal or secondary diagnosis) recorded in the HCUP data [22].

**Figure 4:**
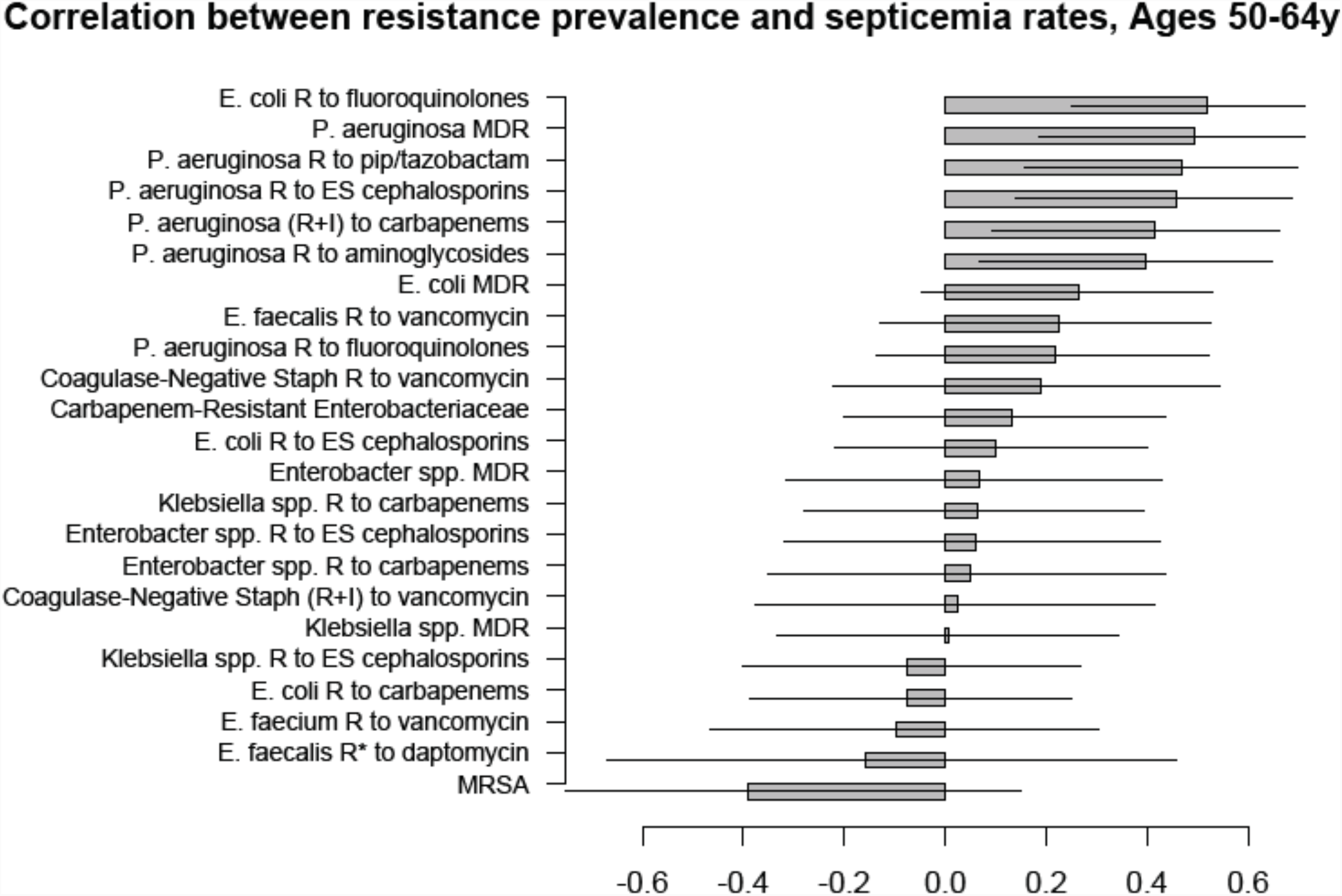
Correlation between state-specific prevalence (percentages) of resistance for different combinations of antibiotics/bacteria in CAUTI samples from hospitalized adults aged 19-64y in the CDC AR Atlas data [21] between 2011-14 and state-specific average annual rates per 100,000 individuals aged 50-64y of septicemia hospitalizations (principal or secondary diagnosis) recorded in the HCUP data [22].

**Figure 5:**
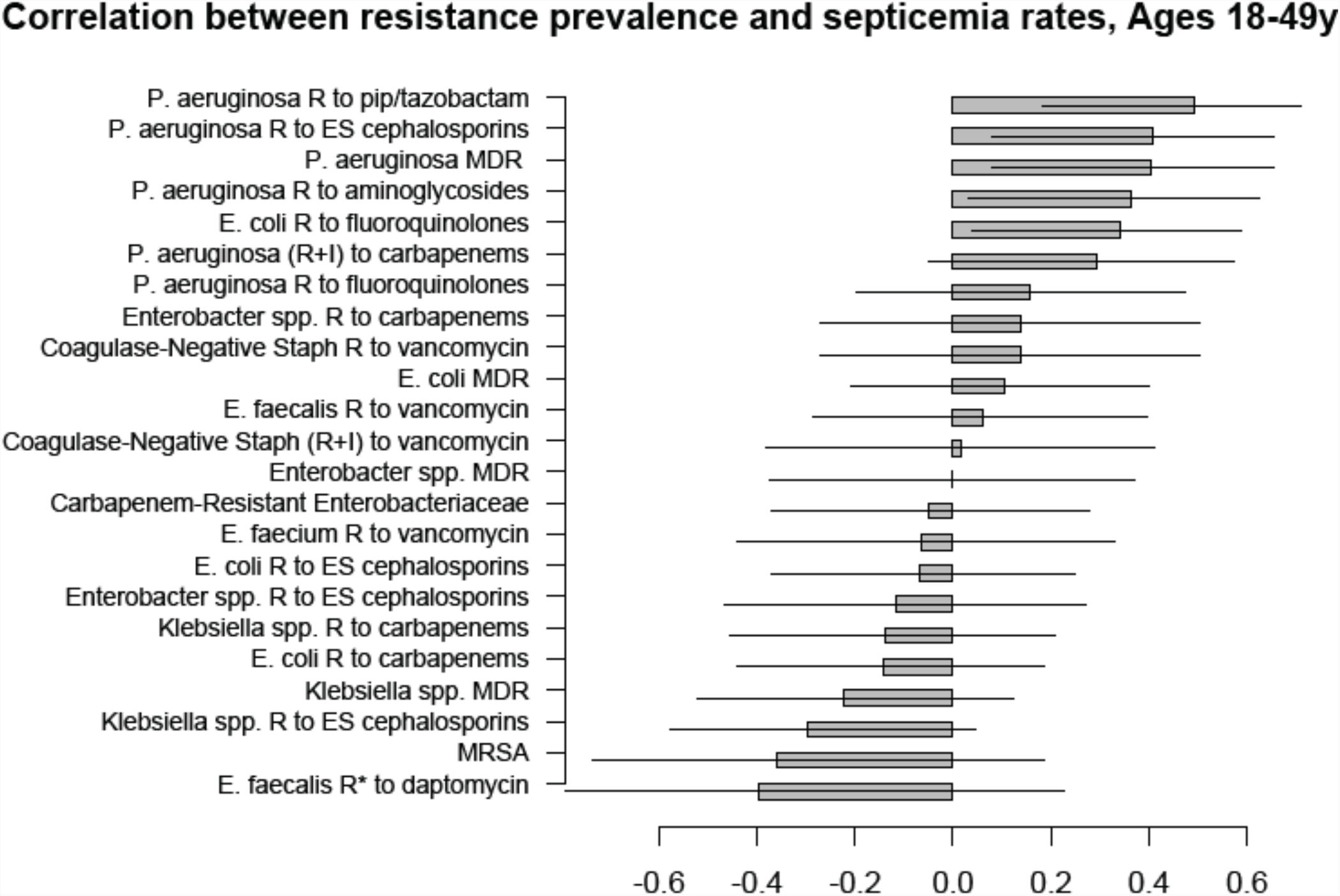
Correlation between state-specific prevalence (percentages) of resistance for different combinations of antibiotics/bacteria in CAUTI samples from hospitalized adults aged 19-64y in the CDC AR Atlas data [21] between 2011-14 and state-specific average annual rates per 100,000 individuals aged 18-49y of septicemia hospitalizations (principal or secondary diagnosis) recorded in the HCUP data [22].

**Figure 6:**
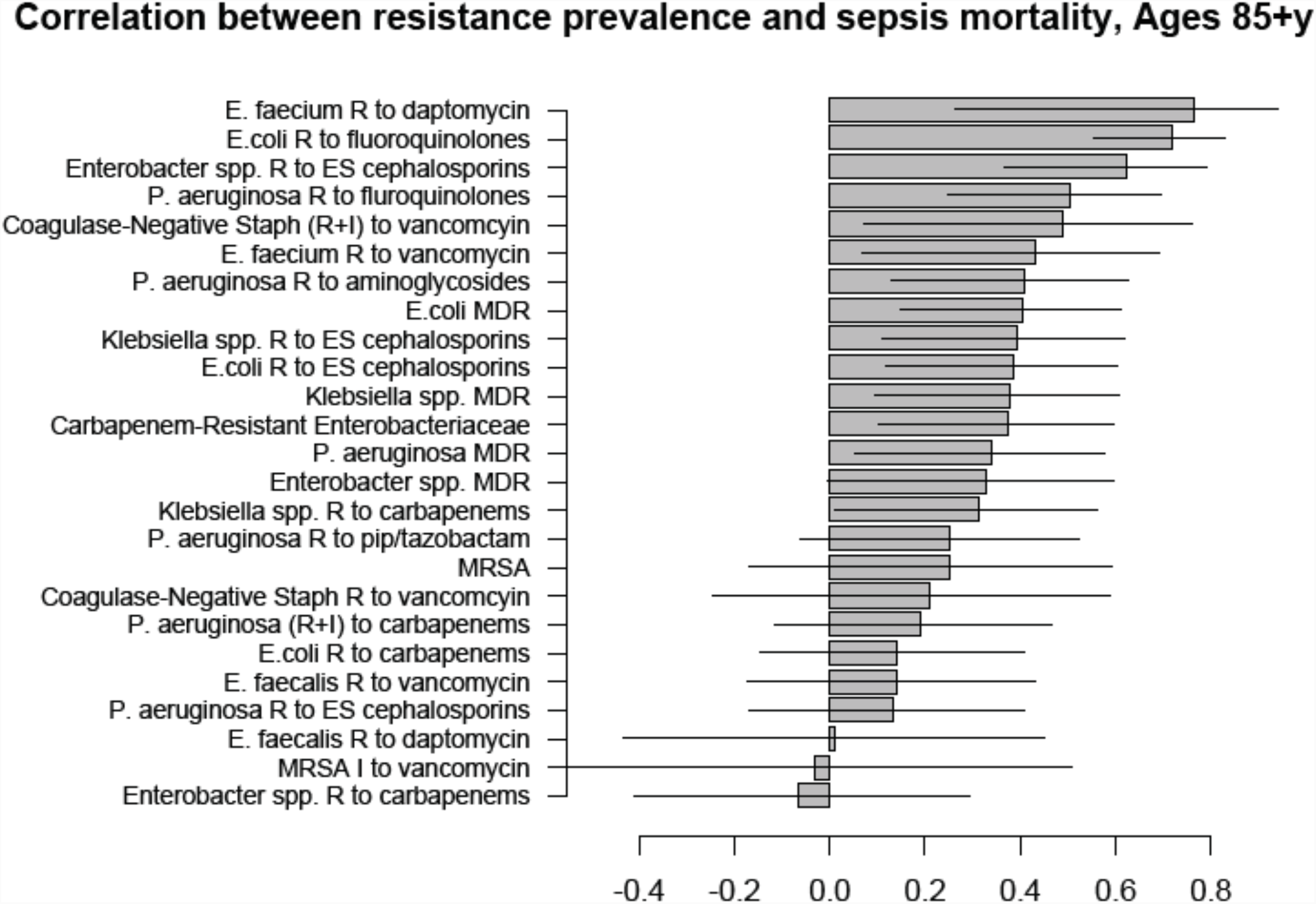
Correlation between state-specific prevalence (percentages) of resistance for different combinations of antibiotics/bacteria in CAUTI samples from hospitalized adults aged 65+y in the CDC AR Atlas data [21] between 2011-14 and state-specific average annual rates per 100,000 individuals aged 85+y of mortality with sepsis listed on the death certificate between 2013-14 [23].

**Figure 7:**
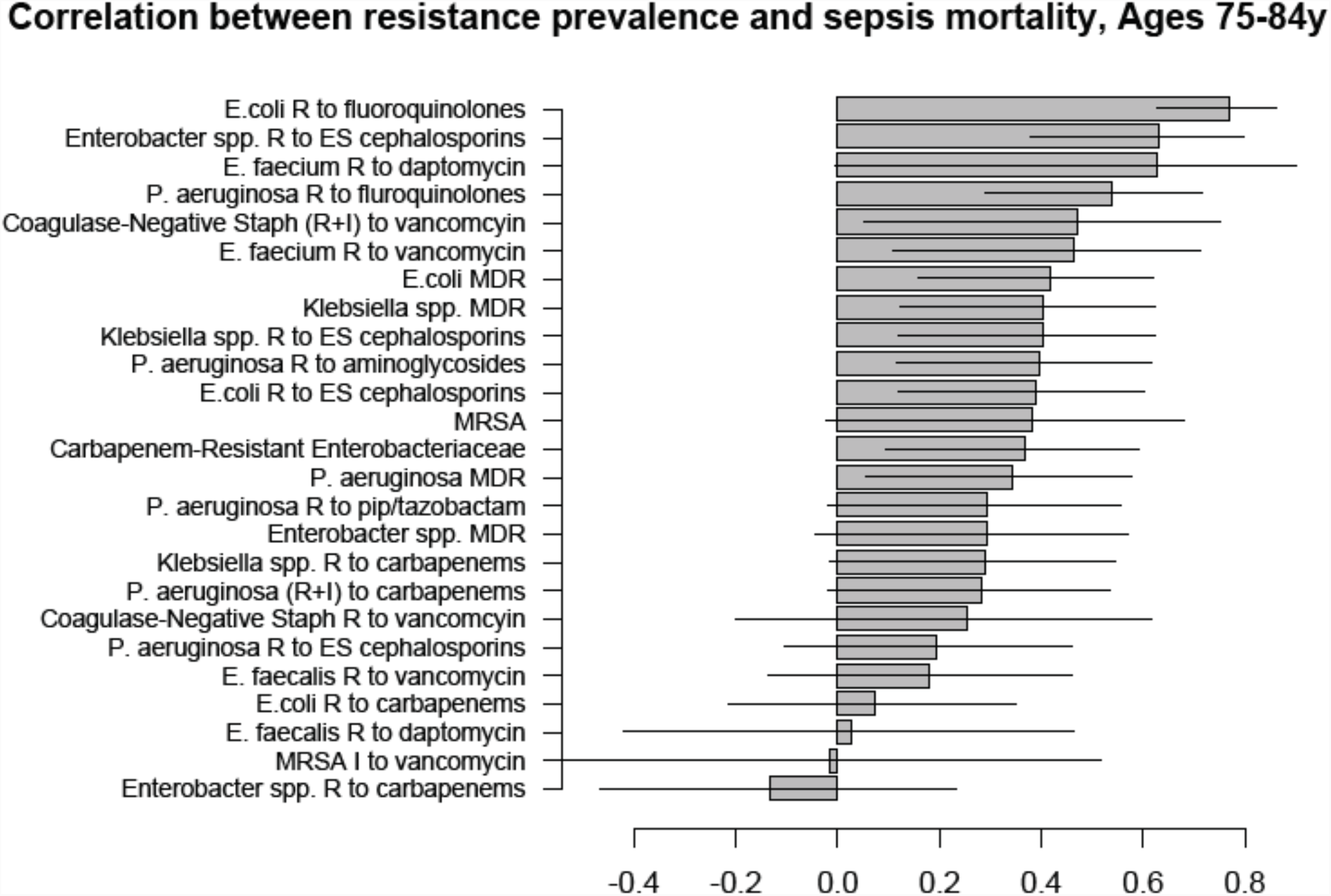
Correlation between state-specific prevalence (percentages) of resistance for different combinations of antibiotics/bacteria in CAUTI samples from hospitalized adults aged 65+y in the CDC AR Atlas data [21] between 2011-14 and state-specific average annual rates per 100,000 individuals aged 75-84y of mortality with sepsis listed on the death certificate between 2013-14 [23].

**Figure 8:**
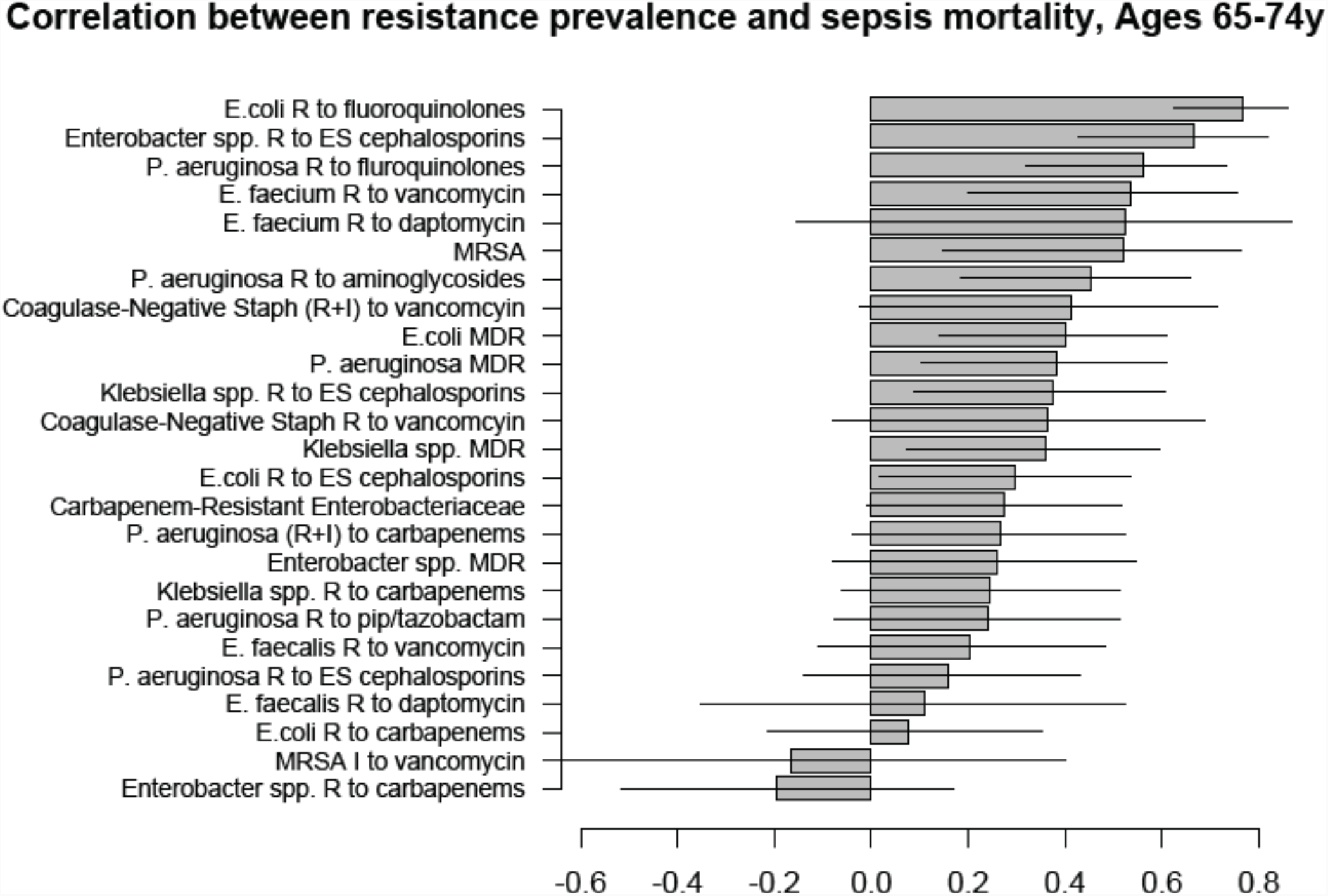
Correlation between state-specific prevalence (percentages) of resistance for different combinations of antibiotics/bacteria in CAUTI samples from hospitalized adults aged 65+y in the CDC AR Atlas data [21] between 2011-14 and state-specific average annual rates per 100,000 individuals aged 65-74y of mortality with sepsis listed on the death certificate between 2013-14 [23].

**Figure 9:**
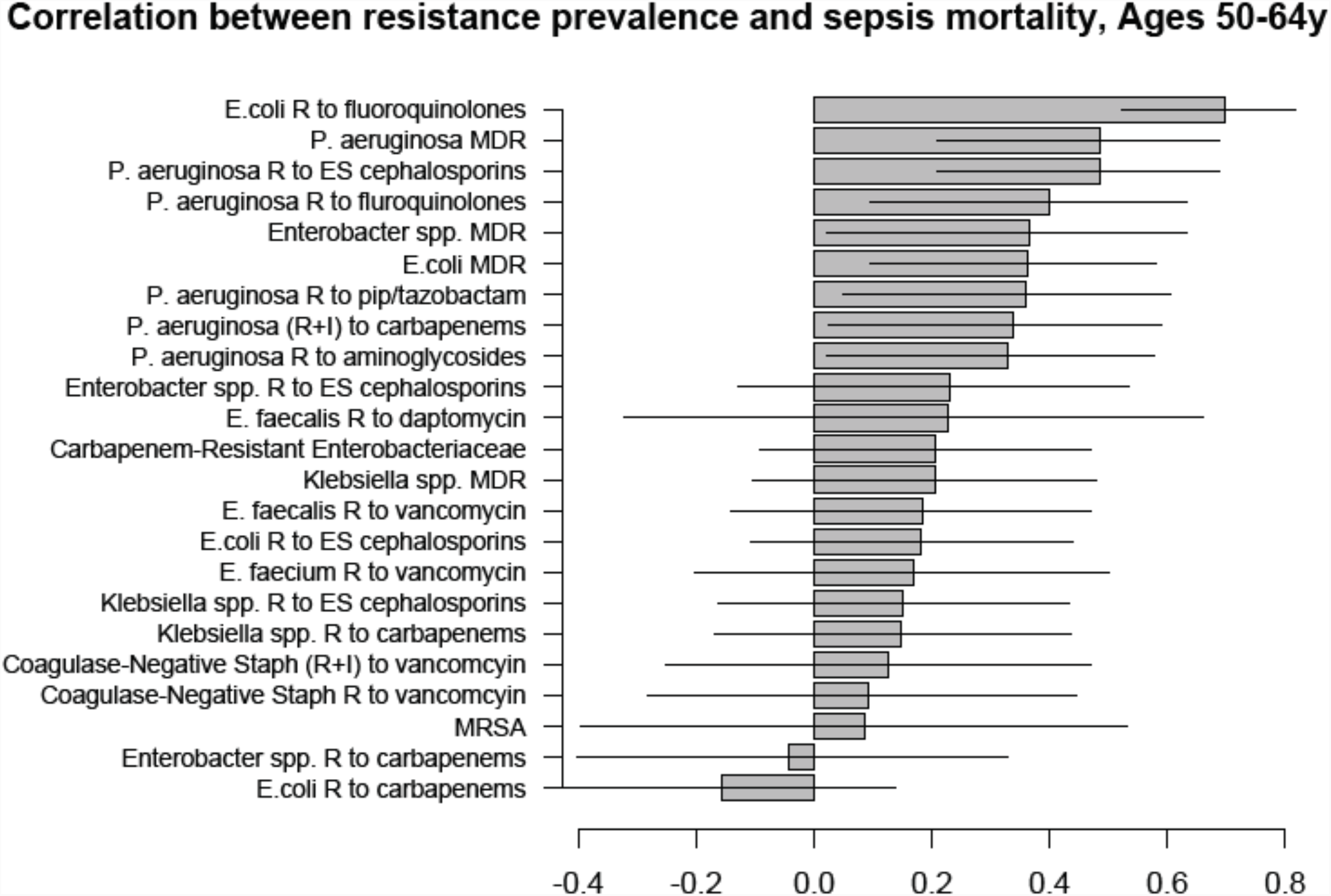
Correlation between state-specific prevalence (percentages) of resistance for different combinations of antibiotics/bacteria in CAUTI samples from hospitalized adults aged 19-64y in the CDC AR Atlas data [21] between 2011-14 and state-specific average annual rates per 100,000 individuals aged 50-64y of mortality with sepsis listed on the death certificate between 2013-14 [23].

**Figure 10:**
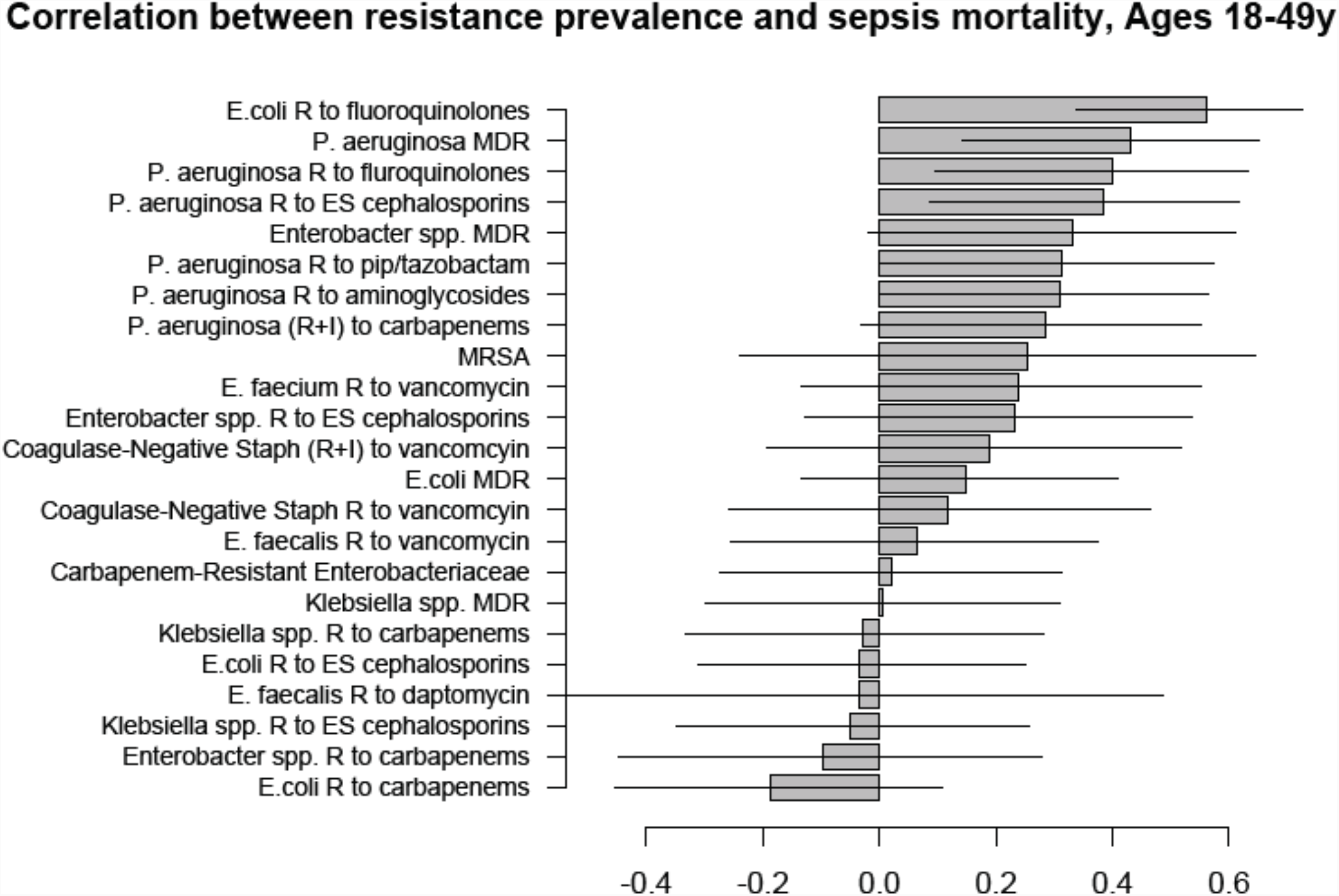
Correlation between state-specific prevalence (percentages) of resistance for different combinations of antibiotics/bacteria in CAUTI samples from hospitalized adults aged 19-64y in the CDC AR Atlas data [21] between 2011-14 and state-specific average annual rates per 100,000 individuals aged 18-49y of mortality with sepsis listed on the death certificate between 2013-14 [23].

A large number of positive correlations between prevalence of antibiotic resistance and rates of septicemia hospitalization/sepsis mortality in different age groups of adults were found, particularly for older adults (Figures 1-10 and Supporting Information), with 66/69 point estimates for the studied correlations for the hospitalization data and 69/75 point estimates for the studied correlations for the mortality data in adults aged over 65y being positive (Figures 1-10). While 99 of the positive estimates for the studied correlations reached nominal statistical significance (48 for hospitalizations and 51 for mortality, uncorrected for multiple comparisons), none of the negative estimates for the studied correlatons did so.

Among the 23-25 combinations of antibiotics/bacteria in the CDC Atlas data [21] included in the analyses for the different age groups/sepsis-related outcomes (septicemia hospitalization or sepsis mortality), prevalence of resistance to fluoroquinolones in *E. coli* had the highest correlation with septicemia hospitalization rates in all age groups over 50y, and the highest correlation with sepsis mortality in all age groups under 85y (Figures 1-10). Additionally, *E. coli* is a major source of Gram-negative septicemia in the US [1], and prevalence of resistance to fluoroquinolones in *E. coli* isolates in both urinary tract and bloodstream infections is high in the US [17,26,27]. We also note that no data on prevalence of resistance to penicillins (save for methicillin) are available in [21]. Figure 11 plots the state-specific prevalence of resistance to fluoroquinolones in E*. coli* isolated in the CAUTI samples in the CDC AR Atlas data [21] in hospitalized adults aged 65+ years, 2011-2014 vs. state-specific average annual rates per 100,000 individuals aged 85+y of hospitalizations with septicemia in either the principal or secondary diagnosis recorded in the Healthcare Cost and Utilization Project (HCUP) database [22], 2011-2012 for the 42 states included in our analyses (Methods). Figure 12 plots the state-specific prevalence of resistance to fluoroquinolones in E*. coli* isolated in the CAUTI samples in individuals aged 65+y vs. rates of sepsis mortality in individuals aged 75-84y. Figures 11,12 suggest strong correlation between prevalence of resitance and rates of septicemia/sepsis-related outcomes (Cor=0.70, 95% CI(0.50,0.83) for hospitalizations; Cor=0.77(0.63,0.86) for mortality). We note that there may be substantial differences in diagnostic practices for septicemia and coding practices for sepsis mortality between states [25]. Nonetheless, Figures 11,12 suggest that there are significant differences in the state-specific septicemia hospitalization and sepsis mortality rates, as well as in the types of states that have low septicemia hospitalization/sepsis mortality rates compared to states that have high septicemia hospitalization/sepsis mortality rates (Discussion and Supporting Information).

**Figure 11:**
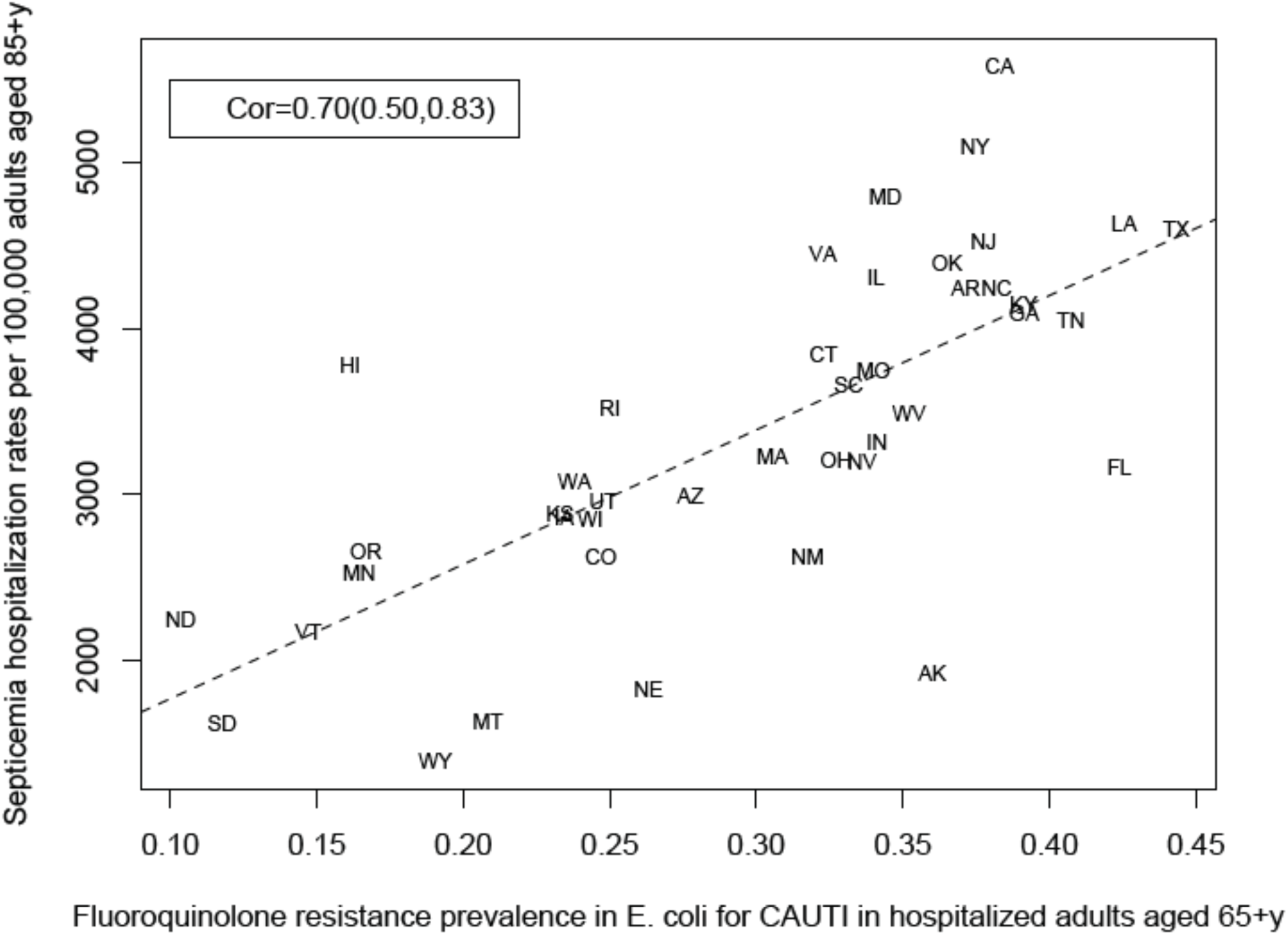
State-specific prevalence (proportion) of resistance to fluoroquinolones in *E. coli* for CAUTI samples in hospitalized adults aged 65+y in the CDC Antibiotic Resistance Atlas data [21], 2011-14 vs. state-specific average annual rates per 100,000 individuals aged 85+y of septicemia hospitalizations (principal or secondary diagnosis) recorded in the HCUP data [22] for the 42 states reporting septicemia hospitalization data between 2011-2012 (Methods).

**Figure 12:**
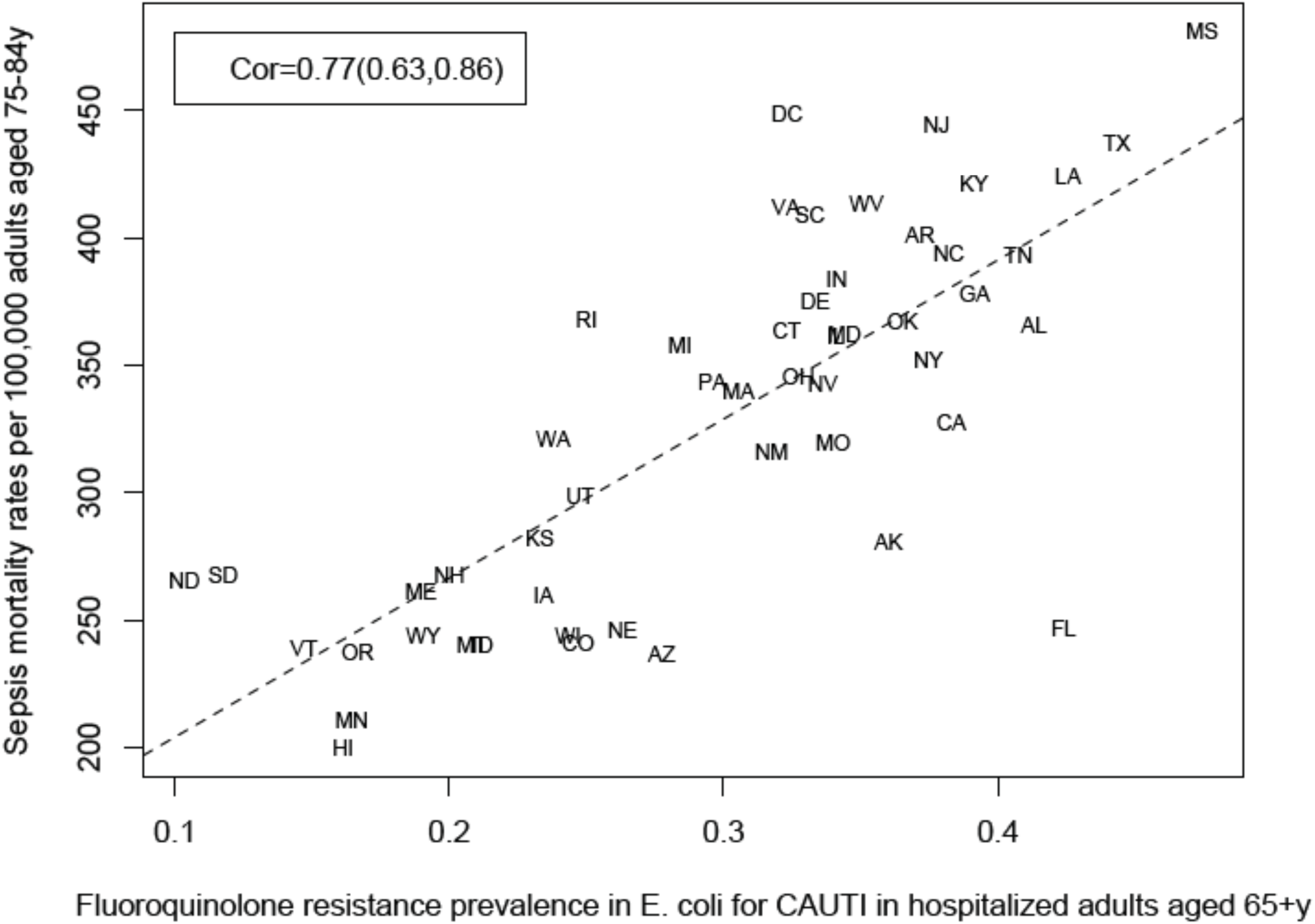
State-specific prevalence (proportion) of resistance to fluoroquinolones in *E. coli* for CAUTI samples in hospitalized adults aged 65+y in the CDC Antibiotic Resistance Atlas data [21], 2011-14 vs. state-specific average annual rates per 100,000 individuals aged 75-84y of sepsis mortality (underlying or contributing cause of death) recorded in the CDC Wonder data [23], 2013-14 for the 50 states + District of Columbia.

We note that there is uncertainty regarding the causal links contributing to several of the correlations presented in Figures 1-10 – see Discussion, as well as the Supporting Information. Finer data are needed to better understand the causal nature of the correlations described in this paper.

## 4. Discussion

Rates of hospitalization with septicemia and sepsis in the diagnosis, and related mortality have risen significantly in last decades in the US [1-3,8], and that rise could not be fully explained by changes in diagnostic practices for septicemia/sepsis [7,8]. Moreover, septicemia hospitalization rates and sepsis mortality rates vary a good deal by state (Supporting Information and [19,20]). Antimicrobial resistance is one of the factors that affects the rates of sepsis-related outcomes, particularly through lack of clearance of resistant infections following antibiotic treatment, with those infections subsequently progressing to septicemia/sepsis and lethal outcomes [9-11]. In this study, we examine the relation between prevalence of antibiotic resistance and rates of septicemia hospitalization as well as sepsis mortality in the US context. For several age groups of adults, and a number of combinations of bacteria/antibiotics, we evaluated the correlations between the state-specific prevalence of resistance in catheter-associated urinary tract infection (CAUTI) samples in the CDC Antibiotic Resistance Safety Atlas data [21] and (a) state-specific rates of hospitalizations with septicemia present in the discharge diagnosis in the HealthCare Cost and Utilization Project (HCUP) data [22]; (b) state-specific rates of mortality with sepsis recorded in the CDC Wonder data [23]. Our results suggest that the state-specific prevalence of resistance to several antibiotics in different bacteria is positively correlated with the rates of hospitalization with septicemia and mortality with sepsis in multiple age groups of adults. Among the different combinations of bacteria/antibiotics in the CDC AR Safety Atlas data [21], prevalence of fluoroquinolone resistance in *E. coli* had the strongest correlation with the rates of hospitalization with septicemia in all age groups over 50y, and with the rates of mortality with sepsis in all age groups 18-84y. Additionally, *E. coli* is the most common source of septicemia among the bacteria covered by the CDC AR Atlas data [1], and prevalence of resistance to fluoroquinolones in *E. coli* isolates in both urinary tract and bloodstream infections is high in the US [17,26,27]. We also note that no data on resistance to penicillins other than methicillin are available in [21], with the use of penicillins found to to be associated with the rates of sepsis in older adults in our related work [19,20]. The results of this study, as well as [19,20] support the association between resistance to/use of certain antibiotics and rates of sepsis-related outcomes, with further work needed to examine the potential utility of replacing certain antibiotics in the treatment of different syndromes on the rates of severe outcomes associated with bacterial infections, including septicemia/sepsis and related mortality.

A possible contributing factor to the various positive correlations that we found (Figures 1-10) is the association between state-specific prevalence of resistance for different pairs of combinations of antibiotics/bacteria. Section S3 of the Supporting Information presents those associations for a number of pairs of combinations of antibiotics/bacteria covered by the CDC Antibiotic Resistance Patient Safety Atlas [21]. Prevalence of resistance for certain combinations of antibiotics/bacteria not covered by the US CDC Atlas data [21] could have also contributed to the correlations found in our analysis. For example, for *E. coli*-associated UTIs recorded in Veterans Affairs data [26], prevalence of resistance to ampicillin/amoxicillin was higher than prevalence of resistance to fluoroquinolones, and prevalence of resistance to trimethoprim/sulfamethoxazole was also high. Additionally, significant associations between presence of resistance to different antibiotics in the samples in [26,17] were found. Thus the correlation between the prevalence of fluoroquinolone resistance in *E. coli* and rates of septicemia hospitalization found in this paper could have been affected not only by the contribution of fluoroquinolone resistance in *E. coli* to the rates of septicemia hospitalization, but also by (i) the contribution of resistance to other antibiotics in *E. coli*, including amoxicillin and trimethoprim/sulfamethoxazole, (ii) the contribution of resistance to fluoroquinolones, and possibly other antibiotics in different bacteria that cause septicemia hospitalization. The overall conclusion supported by our findings is that resistance to different antibiotics in different bacteria (including drug resistance in *E. coli*) contributes to the rates of hospitalization for septicemia/sepsis mortality in the US, which resonates with the findings in [9-11,19,20].

The findings in this paper, as well as other work, e.g. [9-11,19,20] lead to the question whether antibiotic stewardship and replacement of certain antibiotics by others could bring about a reduction in the rates of severe bacterial infections, including sepsis. Some evidence to that effect is provided by the reduction in fluoroquinolone and cephalosporin prescribing in the UK after 2006. Fluoroquinolone use and resistance was found to be associated with MRSA acquisition [14,15]. The drop in the rates of MRSA bacteremia in England between 2006-2011 was sharper than the decline in the rates of MRSA invasive disease in the US during the same period, particularly for the community-associated infections (compare Figures 1 and 3 in [28] with [29]). We also note the high prevalence of fluoroquinolone (levofloxacin) non-susceptibility in community-associated MRSA isolates in the US in the recent years [30]. For severe bacterial infections other than sepsis, reduction in fluoroquinolone and cephalosporin prescribing in England between 2006-2013 was associated with about 75% reduction in the rates of *C. difficile* infections (and even greater reduction in the rates of *C. difficile*-associated deaths) [18,28,31]. While rates of *C. difficile* infection in the US are much higher than in England [32], US rates of *C. difficile* infection have been decreasing during the recent years [33], concomitant with the ongoing decrease in both the inpatient fluoroquinolone use and the (fluoroquinolone-resistant) *C. difficile* epidemic strain NAP1/027 among persons aged ≥65 years [33]. Finally, we note that levels of *E. coli* and *Klebsiella*-associated bacteremia were continuing to rise in England after 2006 while reduction in fluoroquinolone and cephalosporin use was taking place [31,34]. Amoxicillin-clavulanate (AMC, or co-amoxiclav) prescribing in England increased significantly between 2006-2011 [35], and incidence of bacteremia with *E. coli* strains resistant to co-amoxiclav began to increase rapidly after 2006 ([36], Figure 4), with co-amoxiclav resistance in *E. coli*-associated bacteremia exceeding 40% in 2014 [37]. Furthermore, prevalence of co-amoxiclav resistance in *E. coli*-associated bacteremia in England is more than twice as high as the prevalence of co-amoxiclav resistance in *E. coli*-associated UTIs [37], which is also suggestive of the role of co-amoxiclav resistance in progressing to the more severe outcomes resulting from *E. coli* infections. In 2017, the UK has updated its prescribing guidelines for urinary tract infections (UTIs), with nitrofurantoin generally recommended as the first-line option ([37], p. 6). Recently, US FDA has recommended the restriction of fluoroquinolone use for certain conditions (such as uncomplicated UTIs) due to potential adverse effects [38]. At the same time, no recommendations for antibiotics serving as replacement of fluoroquinolones were presented in the FDA guidelines [38]. Such recommendations are needed to optimize the effect of those guidelines on the rates of severe outcomes associated with bacterial infections. For example, data for *E. coli*-associated UTIs in the US [26] suggest that prevalence of resistance varies significantly by antibiotic type, with the highest prevalence of resistance in [26] being for amoxicillin, ampicillin/beta-lactamese inhibitor and fluoroquinolones, with prevalence of resistance in [26] being notably lower for narrow spectrum antibiotics such as nitrofurantoin.

Our study has some limitations. Various confounders could have affected the correlations found in our paper, including the fact that in certain places doctors may be more likely to both assign a diagnosis of sepsis and prescribe antibiotics, particularly in the inpatient setting, creating an association between levels of antibiotic resistance and rates of sepsis. We note that our related work [19,20] suggests associations between outpatient use of certain antibiotics and rates of septicemia hospitalization and sepsis mortality, with those associations presumably being less affected by the practices for assigning a septicemia/sepsis diagnosis and inpatient antibiotic administration. The data in [21] do not contain information on several bacteria that are important sources of septicemia (e.g. MSSA and Streptococci), and resistance to several types of antibiotics (e.g. macrolides, or penicillins other than methicillin). While the HCUP data analyzed in our study generally cover about 97% of all community hospitalizations in the US [22], state-specific variability in the proportion of septicemia hospitalizations that are covered by the HCUP data is possible. Additionally, diagnostic practices for septicemia vary by state. Moreover, we studied correlations between septicemia hospitalization rates, as well as sepsis mortality rates in various subgroups of the elderly (e.g. aged 74-85y) and non-elderly adults and prevalence of antibiotic resistance in CAUTI samples from elderly (aged 65+y) or non-elderly (aged 19-64y) hospitalized adult patients correspondingly. Finally, data on antimicrobial resistance were available for the 2011-2014 period [21], with low counts/missing data for some combinations of bacteria/antibiotics in certain states during certain years; data on septicemia hospitalizations were available through the end of 2012; we chose the 2013-2014 period to evaluate the average annual state-specific sepsis mortality rates due to temporal changes in coding practices for sepsis on death certificates [25]. Correspondingly, we studied correlations between the state-specific prevalence of antibiotic resistance between 2011-2014 and (a) septicemia hospitalization rates between 2011-2012; (b) sepsis mortality rates between 2013-2014. We expect that those sources of noise/incompatibility should generally bias the correlation estimates towards the null, reducing precision rather than creating spurious associations.

We believe that despite certain limitations, our paper suggests a possibly important effect of the prevalence of antibiotic resistance on the rates of septicemia hospitalization and sepsis mortality in US adults. Our results support the need for (i) examing the utility of replacement of certain antibiotics by certain others in the treatment of different syndromes for reducing the rates of severe outcomes associated with bacterial infections; (ii) enhancing antibiotic stewardship (including the use of fluoroquinolones), both in the inpatient and the outpatient settings; (iii) stepping up efforts for preventing acquisition of antibiotic-resistant bacteria (particularly *E. coli* in the elderly) [39]. Additionally, the relation between antibiotic resistance as well as antibiotic use and the rates of severe bacterial infections, including sepsis, as well as the associated mortality outcomes [9-12,19,20], and the limited options for antibiotic replacement support the need for new antibiotics. This need may not be fulfilled through the current system of antibiotic development/production, and additional incentives for antibacterial research and development are worth considering [40].

## Acknowledgement

We would like to thank the HCUP Partner states that voluntarily provide their data to the project, and without whom this research would not be possible (https://www.hcup-us.ahrq.gov/partners.jsp).

