## Supporting Information for "Antimicrobial resistance prevalence, rates of hospitalization with septicemia and rates of mortality with sepsis in adults in different US states"

Supporting Information for "**Antimicrobial resistance prevalence and rates of hospitalization with septicemia in the diagnosis in adults in different US states**"

**Section S1**: Rates hospitalization with septicemia in different US states

Tables S1-S5 show the 10 states with the highest average annual rates of hospitalizations with septicemia (ICD-9 codes 038.xx present in either the principal or secondary discharge diagnosis) recorded in the HealthCare Cost and Utilization Project (HCUP) database [1] between 2011-12 per 100,000 individuals aged 85+y (Table S1), aged 75-84y (Table S2), aged 65-74y (Table S3), aged 50-64y (Table S4), and aged 18-49y (Table S5), as well as the 10 states with the lowest septicemia hospitalization rates in the corresponding age group among the 42 states that reported septicemia hospitalizations to HCUP between 2011-12. Those 42 states are: Alaska, Arkansas, Arizona, California, Colorado, Connecticut, Florida, Georgia, Hawaii, Iowa, Illinois, Indiana, Kansas, Kentucky, Louisiana, Massachusetts, Maryland, Minnesota, Missouri, Montana, North Carolina, North Dakota, Nebraska, New Jersey, New Mexico, Nevada, New York, Ohio, Oklahoma, Oregon, Rhode Island, South Carolina, South Dakota, Tennessee, Texas, Utah, Virginia, Vermont, Washington, Wisconsin, West Virginia, Wyoming.

We note that the rates in Tables S1-S5 may reflect significant differences in diagnostic practices between some states. We also note that there is a degree of consistency between Tables S1-S5, particularly in terms of the states with the lowest rates of hospitalizations with septicemia. Those states with the lowest septicemia hospitalization rates tend to be northern states with low population density, which may be suggestive of the differences in the rates of bacterial transmission between different states.

| State | Septicemia hospitalization rate | | State | Septicemia hospitalization rate |
|---|---|---|---|---|
| California | 5557 | | Wyoming | 1564 |
| New York | 5022 | | Montana | 1723 |
| Maryland | 4732 | | South Dakota | 1789 |
| Louisiana | 4621 | | Nebraska | 1946 |
| Virginia | 4542 | | Alaska | 2008 |
| Texas | 4485 | | Vermont | 2215 |
| New Jersey | 4437 | | North Dakota | 2231 |
| Oklahoma | 4340 | | Minnesota | 2501 |
| Illinois | 4312 | | New Mexico | 2604 |
| North Carolina | 4272 | | Colorado | 2699 |

**Table S1:** Average annual rates per 100,000 individuals aged 85+y of septicemia hospitalizations (principal or secondary diagnosis) recorded in the HealthCare Cost and Utilization Project (HCUP) data [1] for the states with the highest 10 rates and the lowest 10 rates among the 42 states reporting between 2011-2012 (Methods).

| State | Septicemia hospitalization rate | | State | Septicemia hospitalization rate |
|---|---|---|---|---|
| California | 3047 | | Wyoming | 996 |
| Maryland | 2950 | | Montana | 1255 |
| New York | 2708 | | Alaska | 1259 |
| Texas | 2700 | | South Dakota | 1471 |
| Louisiana | 2652 | | Nebraska | 1493 |
| Kentucky | 2632 | | Vermont | 1506 |
| New Jersey | 2615 | | Minnesota | 1706 |
| Illinois | 2594 | | New Mexico | 1759 |
| North Carolina | 2587 | | Colorado | 1813 |
| Virginia | 2550 | | Wisconsin | 1859 |

**Table S2:** Average annual rates per 100,000 individuals aged 75-84y of septicemia hospitalizations (principal or secondary diagnosis) recorded in the HealthCare Cost and Utilization Project (HCUP) data [1] for the states with the highest 10 rates and the lowest 10 rates among the 42 states reporting between 2011-2012 (Methods).

| State | Septicemia hospitalization rate | | State | Septicemia hospitalization rate |
|---|---|---|---|---|
| Maryland | 1547 | | Wyoming | 606 |
| Missouri | 1514 | | Vermont | 686 |
| California | 1504 | | Montana | 737 |
| Texas | 1469 | | Alaska | 752 |
| Kentucky | 1461 | | South Dakota | 825 |
| Tennessee | 1457 | | Nebraska | 908 |
| Oklahoma | 1445 | | Minnesota | 985 |
| Louisiana | 1432 | | New Mexico | 986 |
| North Carolina | 1416 | | Colorado | 996 |
| Illinois | 1381 | | Oregon | 1037 |

**Table S3:** Average annual rates per 100,000 individuals aged 65-74y of septicemia hospitalizations (principal or secondary diagnosis) recorded in the HealthCare Cost and Utilization Project (HCUP) data [1] for the states with the highest 10 rates and the lowest 10 rates among the 42 states reporting between 2011-2012 (Methods).

| State | Septicemia hospitalization rate | | State | Septicemia hospitalization rate |
|---|---|---|---|---|
| Tennessee | 720 | | Wyoming | 258 |
| Missouri | 719 | | Vermont | 301 |
| Maryland | 708 | | Alaska | 311 |
| Oklahoma | 693 | | Montana | 347 |
| Arizona | 693 | | Minnesota | 415 |
| West Virginia | 687 | | Nebraska | 416 |
| Kentucky | 684 | | Connecticut | 434 |
| Louisiana | 665 | | South Dakota | 436 |
| North Carolina | 655 | | Massachusetts | 439 |
| Texas | 650 | | Iowa | 446 |

**Table S4:** Average annual rates per 100,000 individuals aged 50-64y of septicemia hospitalizations (principal or secondary diagnosis) recorded in the HealthCare Cost and Utilization Project (HCUP) data [1] for the states with the highest 10 rates and the lowest 10 rates among the 42 states reporting between 2011-2012 (Methods).

| State | Septicemia hospitalization rate | | State | Septicemia hospitalization rate |
|---|---|---|---|---|
| Arizona | 255 | | Wyoming | 80 |
| West Virginia | 254 | | Vermont | 86 |
| Tennessee | 237 | | Nebraska | 103 |
| Maryland | 237 | | Alaska | 105 |
| Missouri | 225 | | South Dakota | 117 |
| New Mexico | 210 | | Minnesota | 120 |
| Kentucky | 206 | | Massachusetts | 120 |
| Hawaii | 200 | | Connecticut | 121 |
| Ohio | 195 | | Montana | 122 |
| Oklahoma | 194 | | Iowa | 122 |

**Table S5:** Average annual rates per 100,000 individuals aged 18-49y of septicemia hospitalizations (principal or secondary diagnosis) recorded in the HealthCare Cost and Utilization Project (HCUP) data [1] for the states with the highest 10 rates and the lowest 10 rates among the 42 states reporting between 2011-2012 (Methods).

**Section S2**: Rates of mortality with sepsis in different US states

Tables S6-S10 show the 10 states with the highest average annual rates of mortality with sepsis (ICD-10 codes A40-41.xx present as either contributing or underlying causes of death in [2]) between 2013-14 per 100,000 individuals aged 85+y (Table S6), aged 75-84y (Table S7), aged 65-74y (Table S8), aged 50-64y (Table S9), and aged 18-49y (Table S10), as well as the 10 states with the lowest sepsis mortality rates in the corresponding age group. States with the higher sepsis mortality rates tend to be in the South/East, while states with lower sepsis mortality rates tend to be in the North/West.

| State | Sepsis mortality rate | | State | Sepsis mortality rate |
|---|---|---|---|---|
| Mississippi | 1114.6 | | Minnesota | 418.4 |
| New Jersey | 1043.9 | | Hawaii | 461.9 |
| Virginia | 987.9 | | Nebraska | 465.6 |
| Texas | 983.6 | | South Dakota | 489.8 |
| Louisiana | 977 | | Wisconsin | 516.6 |
| District of Columbia | 964.7 | | Oregon | 519.2 |
| Arkansas | 920.7 | | Arizona | 519.3 |
| North Carolina | 916.3 | | Montana | 519.4 |
| West Virginia | 895.7 | | Florida | 542.8 |
| Georgia | 875.7 | | Colorado | 545 |

**Table S6:** Average annual rates per 100,000 individuals aged 85+y of sepsis mortality (ICD-10 codes A40-41.xx present as either contributing or underlying causes of death in [2]) for the states with the highest 10 rates and the lowest 10 rates between 2013-2014.

| State | Sepsis mortality rate | | State | Sepsis mortality rate |
|---|---|---|---|---|
| Mississippi | 478.4 | | Minnesota | 205.6 |
| New Jersey | 456.1 | | Montana | 231.6 |
| District of Columbia | 438.5 | | Arizona | 233.2 |
| Texas | 430 | | Wisconsin | 235 |
| Virginia | 418.2 | | South Dakota | 235.3 |
| Louisiana | 411.8 | | Hawaii | 235.4 |
| West Virginia | 410 | | Colorado | 235.6 |
| Arkansas | 408.1 | | Wyoming | 237.8 |
| South Carolina | 398.8 | | Nebraska | 245.2 |
| Kentucky | 393.8 | | Oregon | 247.1 |

**Table S7:** Average annual rates per 100,000 individuals aged 75-84y of sepsis mortality (ICD-10 codes A40-41.xx present as either contributing or underlying causes of death in [2]) for the states with the highest 10 rates and the lowest 10 rates between 2013-2014.

| State | Sepsis mortality rate | | State | Sepsis mortality rate |
|---|---|---|---|---|
| Mississippi | 224.4 | | Alaska | 82.2 |
| Louisiana | 192 | | Montana | 82.2 |
| Texas | 187.9 | | South Dakota | 83.9 |
| Kentucky | 181.6 | | Minnesota | 84.8 |
| West Virginia | 181.1 | | Wisconsin | 91.4 |
| Arkansas | 178.9 | | Hawaii | 94.6 |
| Alabama | 177.7 | | Colorado | 95.3 |
| Tennessee | 174.6 | | Oregon | 95.5 |
| District of Columbia | 173.1 | | Nebraska | 97.9 |
| Indiana | 173 | | Iowa | 103.9 |

**Table S8:** Average annual rates per 100,000 individuals aged 65-74y of sepsis mortality (ICD-10 codes A40-41.xx present as either contributing or underlying causes of death in [2]) for the states with the highest 10 rates and the lowest 10 rates between 2013-2014.

| State | Sepsis mortality rate | | State | Sepsis mortality rate |
|---|---|---|---|---|
| Mississippi | 92.2 | | Minnesota | 29 |
| Alabama | 80.3 | | Vermont | 30.1 |
| Oklahoma | 78 | | Wisconsin | 31.2 |
| West Virginia | 75.7 | | New Hampshire | 31.3 |
| Louisiana | 75.1 | | Hawaii | 32.3 |
| Kentucky | 74.9 | | Colorado | 34.1 |
| Arkansas | 73.4 | | Alaska | 34.8 |
| Texas | 70.4 | | Nebraska | 35.9 |
| Tennessee | 69.9 | | Utah | 36.4 |
| South Carolina | 69.7 | | Maine | 37 |

**Table S9:** Average annual rates per 100,000 individuals aged 50-64y of sepsis mortality (ICD-10 codes A40-41.xx present as either contributing or underlying causes of death in [2]) for the states with the highest 10 rates and the lowest 10 rates between 2013-2014.

| State | Sepsis mortality rate | | State | Sepsis mortality rate |
|---|---|---|---|---|
| Mississippi | 15.2 | | Vermont | 3.5 |
| West Virginia | 15.2 | | Minnesota | 3.6 |
| Alabama | 13.5 | | Iowa | 4 |
| Arkansas | 12.8 | | Wisconsin | 4.7 |
| Kentucky | 12.1 | | Utah | 4.8 |
| Tennessee | 11.4 | | New York | 5 |
| South Carolina | 11.3 | | Colorado | 5.1 |
| Louisiana | 11 | | Alaska | 5.1 |
| Georgia | 10.4 | | New Hampshire | 5.5 |
| Oklahoma | 10 | | Idaho | 5.6 |

**Table S10:** Average annual rates per 100,000 individuals aged 18-49y of sepsis mortality (ICD-10 codes A40-41.xx present as either contributing or underlying causes of death in [2]) for the states with the highest 10 rates and the lowest 10 rates between 2013-2014.

**Section S3:** Correlations between state-specific prevalence of resistance for different combinations of antibiotics/bacteria

Tables S11-S16 exhibit correlations between state-specific prevalence of resistance in catheter-associated urinary tract infection (CAUTI) samples in the CDC Antibiotic Resistance Patient Safety Atlas data [2] between 2011-2014 for certain combinations of antibiotics/bacteria. Tables S11-S13 shows the results for CAUTI samples in hospitalized adults aged 65+y, while Tables S14-S16 show the results for CAUTI samples in hospitalized adults aged 19-64y. A large number of positive correlations between prevalence of resistance for different combinations of bacteria/antibiotics were found, including resistance to different antibiotics in the same bacteria, resistance to the same antibiotic in different bacteria, as well as resistance to different antibiotics in different bacteria. Those correlations may be partly affected by patterns of antibiotic usage in different states (e.g. [4]), including antibiotic usage during urinary catheterization procedures, as well as patterns of cross-resistance to different antibiotics in the same bacteria, e.g. [5]. As suggested in the Discussion, those correlations may contribute to the correlations between prevalence of resistance and rates of hospitalizations with septicemia presented in Tables 1-10 in the main manuscript.

| | E. coli R to fluoroquinolones | E. coli MDR | E. coli R to ES cephalosporins | P. aeruginosa R to fluroquinolones | P. aeruginosa R to aminoglycosides | P. aeruginosaDR | P. aeruginosa R to pip/tazobactam |
|---|---|---|---|---|---|---|---|
| E. coli R to fluoroquinolones | 1 | 0.69 (0.49,0.82) | 0.54 (0.28,0.73) | 0.63 (0.39,0.79) | 0.66 (0.42,0.81) | 0.48 (0.19,0.7) | 0.37 (0.03,0.63) |
| E. coli MDR | 0.69 (0.49,0.82) | 1 | 0.81 (0.66,0.89) | 0.39 (0.07,0.63) | 0.51 (0.22,0.72) | 0.41 (0.1,0.65) | 0.22 (-0.13,0.52) |
| E. coli R to ES cephalosporins | 0.54 (0.28,0.73) | 0.81 (0.66,0.89) | 1 | 0.31 (-0.02,0.57) | 0.4 (0.08,0.64) | 0.36 (0.04,0.61) | 0.25 (-0.1,0.54) |
| P. aeruginosa R to fluroquinolones | 0.63 (0.39,0.79) | 0.39 (0.07,0.63) | 0.31 (-0.02,0.57) | 1 | 0.84 (0.71,0.91) | 0.86 (0.75,0.93) | 0.69 (0.46,0.83) |
| P. aeruginosa R to aminoglycosides | 0.66 (0.42,0.81) | 0.51 (0.22,0.72) | 0.4 (0.08,0.64) | 0.84 (0.71,0.91) | 1 | 0.82 (0.68,0.91) | 0.54 (0.24,0.74) |
| P. aeruginosa MDR | 0.48 (0.19,0.7) | 0.41 (0.1,0.65) | 0.36 (0.04,0.61) | 0.86 (0.75,0.93) | 0.82 (0.68,0.91) | 1 | 0.71 (0.49,0.85) |
| P. aeruginosa R to pip/tazobactam | 0.37 (0.03,0.63) | 0.22 (-0.13,0.52) | 0.25 (-0.1,0.54) | 0.69 (0.46,0.83) | 0.54 (0.24,0.74) | 0.71 (0.49,0.85) | 1 |
| P. aeruginosa R to ES cephalosporins | 0.15 (-0.18,0.45) | 0.11 (-0.22,0.42) | 0.17 (-0.16,0.47) | 0.49 (0.2,0.7) | 0.43 (0.13,0.66) | 0.67 (0.44,0.82) | 0.65 (0.41,0.81) |

**Table S11**: Correlations between state-specific prevalence of resistance in the CDC Antibiotic Resistance Patient Safety Atlas data [3] between 2011-2014 for certain combinations of antibiotics/bacteria in CAUTI samples in adults aged 65+y.

| | Klebsiella spp. MDR | Klebsiella spp. R to carbapenems | Klebsiella spp. R to ES cephalosporins | Carbapenem-R Enterobacteriaceae | E. faecalis R to vancomycin | E. faecium R to vancomycin | Enterobacter spp. R to ES cephalosporins |
|---|---|---|---|---|---|---|---|
| E. coli R to fluoroquinolones | 0.58 (0.32,0.77) | 0.46 (0.16,0.69) | 0.61 (0.35,0.78) | 0.45 (0.15,0.67) | 0.46 (0.15,0.69) | 0.58 (0.23,0.8) | 0.68 (0.43,0.83) |
| E. coli MDR | 0.65 (0.41,0.81) | 0.66 (0.42,0.81) | 0.67 (0.44,0.82) | 0.61 (0.37,0.78) | 0.42 (0.1,0.66) | 0.52 (0.14,0.76) | 0.41 (0.07,0.67) |
| E. coli R to ES cephalosporins | 0.64 (0.39,0.8) | 0.66 (0.43,0.81) | 0.71 (0.5,0.84) | 0.63 (0.39,0.79) | 0.36 (0.03,0.62) | 0.41 (0,0.7) | 0.4 (0.05,0.66) |
| P. aeruginosa R to fluroquinolones | 0.48 (0.18,0.7) | 0.46 (0.15,0.68) | 0.54 (0.25,0.74) | 0.44 (0.13,0.67) | 0.42 (0.11,0.66) | 0.27 (-0.15,0.61) | 0.38 (0.04,0.65) |
| P. aeruginosa R to aminoglycosides | 0.59 (0.33,0.77) | 0.6 (0.34,0.78) | 0.6 (0.34,0.78) | 0.55 (0.27,0.75) | 0.46 (0.14,0.68) | 0.48 (0.09,0.74) | 0.42 (0.08,0.68) |
| P. aeruginosa MDR | 0.54 (0.26,0.74) | 0.59 (0.32,0.77) | 0.58 (0.31,0.76) | 0.59 (0.32,0.77) | 0.42 (0.1,0.66) | 0.09 (-0.33,0.48) | 0.25 (-0.12,0.55) |
| P. aeruginosa R to pip/tazobactam | 0.33 (-0.01,0.61) | 0.37 (0.03,0.64) | 0.37 (0.03,0.63) | 0.38 (0.04,0.64) | 0.23 (-0.13,0.53) | -0.19 (-0.55,0.23) | 0.24 (-0.13,0.55) |

| | | | | | | | |
|---|---|---|---|---|---|---|---|
| P. aeruginosa R to ES cephalosporins | 0.42 (0.1,0.66) | 0.4 (0.09,0.65) | 0.35 (0.03,0.61) | 0.34 (0.01,0.6) | 0.25 (-0.1,0.54) | -0.14 (-0.52,0.28) | 0.16 (-0.2,0.49) |

**Table S12**: Correlations between state-specific prevalence of resistance in the CDC Antibiotic Resistance Patient Safety Atlas data [3] between 2011-2014 for certain combinations of antibiotics/bacteria in CAUTI samples in adults aged 65+y.

| | Klebsiella spp. MDR | Klebsiella spp. R to carbapenems | Klebsiella spp. R to ES cephalosporins | Carbapenem-R Enterobacteriaceae | E. faecalis R to vancomycin | E. faecium R to vancomycin | Enterobacter spp. R to ES cephalosporins |
|---|---|---|---|---|---|---|---|
| Klebsiella spp. MDR | 1 | 0.92 (0.85,0.96) | 0.95 (0.9,0.97) | 0.88 (0.77,0.94) | 0.49 (0.19,0.71) | 0.43 (0.03,0.71) | 0.66 (0.41,0.82) |
| Klebsiella spp. R to carbapenems | 0.92 (0.85,0.96) | 1 | 0.9 (0.81,0.95) | 0.95 (0.91,0.98) | 0.54 (0.25,0.74) | 0.42 (0.02,0.71) | 0.55 (0.24,0.76) |
| Klebsiella spp. R to ES cephalosporins | 0.95 (0.9,0.97) | 0.9 (0.81,0.95) | 1 | 0.88 (0.77,0.94) | 0.54 (0.25,0.74) | 0.44 (0.05,0.72) | 0.68 (0.42,0.83) |
| Carbapenem-R Enterobacteriaceae | 0.88 (0.77,0.94) | 0.95 (0.91,0.98) | 0.88 (0.77,0.94) | 1 | 0.49 (0.18,0.71) | 0.37 (-0.05,0.68) | 0.52 (0.2,0.74) |
| E. faecalis R to vancomycin | 0.49 (0.19,0.71) | 0.54 (0.25,0.74) | 0.54 (0.25,0.74) | 0.49 (0.18,0.71) | 1 | 0.16 (-0.26,0.53) | 0.19 (-0.17,0.51) |
| E. faecium R to vancomycin | 0.43 (0.03,0.71) | 0.42 (0.02,0.71) | 0.44 (0.05,0.72) | 0.37 (-0.05,0.68) | 0.16 (-0.26,0.53) | 1 | 0.63 (0.29,0.83) |
| Enterobacter spp. R to ES cephalosporins | 0.66 (0.41,0.82) | 0.55 (0.24,0.76) | 0.68 (0.42,0.83) | 0.52 (0.2,0.74) | 0.19 (-0.17,0.51) | 0.63 (0.29,0.83) | 1 |

**Table S13**: Correlations between state-specific prevalence of resistance in the CDC Antibiotic Resistance Patient Safety Atlas data [3] between 2011-2014 for certain combinations of antibiotics/bacteria in CAUTI samples in adults aged 65+y.

| | E. coli R to fluoroquinolones | E. coli MDR | E. coli R to ES cephalosporins | P. aeruginosa R to fluroquinolones | P. aeruginosa R to aminoglycosides | P. aeruginosa MDR | P. aeruginosa R to pip/tazobactam | P. aeruginosa R to ES cephalosporins |
|---|---|---|---|---|---|---|---|---|
| E. coli R to fluoroquinolones | 1 | 0.58 (0.33,0.75) | 0.42 (0.13,0.65) | 0.47 (0.15,0.7) | 0.39 (0.05,0.64) | 0.58 (0.3,0.77) | 0.28 (-0.07,0.56) | 0.53 (0.24,0.74) |
| E. coli MDR | 0.58 (0.33,0.75) | 1 | 0.82 (0.69,0.9) | 0.27 (-0.08,0.56) | 0.37 (0.04,0.63) | 0.47 (0.16,0.7) | 0.19(- 0.16,0.49) | 0.45 (0.14,0.69) |
| E. coli R to ES cephalosporins | 0.42 (0.13,0.65) | 0.82 (0.69,0.9) | 1 | 0.16 (-0.19,0.48) | 0.19 (-0.16,0.5) | 0.2 (-0.15,0.5) | -0.14 (-0.46,0.21) | 0.21 (-0.14,0.51) |
| P. aeruginosa R to fluroquinolones | 0.47 (0.15,0.7) | 0.27 (-0.08,0.56) | 0.16 (-0.19,0.48) | 1 | 0.78 (0.6,0.89) | 0.76 (0.57,0.88) | 0.36 (0.02,0.63) | 0.45 (0.13,0.69) |
| P. aeruginosa R to aminoglycosides | 0.39 (0.05,0.64) | 0.37 (0.04,0.63) | 0.19 (-0.16,0.5) | 0.78 (0.6,0.89) | 1 | 0.89 (0.78,0.94) | 0.59 (0.32,0.78) | 0.5 (0.19,0.72) |
| P. aeruginosa MDR | 0.58 (0.3,0.77) | 0.47 (0.16,0.7) | 0.2 (-0.15,0.5) | 0.76 (0.57,0.88) | 0.89 (0.78,0.94) | 1 | 0.67 (0.43,0.82) | 0.71 (0.5,0.85) |
| P. aeruginosa R to pip/tazobactam | 0.28 (-0.07,0.56) | 0.19 (-0.16,0.49) | -0.14 (-0.46,0.21) | 0.36 (0.02,0.63) | 0.59 (0.32,0.78) | 0.67 (0.43,0.82) | 1 | 0.65 (0.4,0.81) |
| P. aeruginosa R to ES cephalosporins | 0.53 (0.24,0.74) | 0.45 (0.14,0.69) | 0.21 (-0.14,0.51) | 0.45 (0.13,0.69) | 0.5 (0.19,0.72) | 0.71 (0.5,0.85) | 0.65 (0.4,0.81) | 1 |
| P. aeruginosa (R+I) to carbapenems | 0.45 (0.13,0.68) | 0.33 (0,0.6) | 0.07 (-0.27,0.4) | 0.67 (0.43,0.83) | 0.78 (0.6,0.88) | 0.9 (0.8,0.95) | 0.56 (0.28,0.76) | 0.63 (0.37,0.8) |

**Table S14**: Correlations between state-specific prevalence of resistance in the CDC Antibiotic Resistance Patient Safety Atlas data [3] between 2011-2014 for certain combinations of antibiotics/bacteria in CAUTI samples in adults aged 19-64y.

| | Klebsiella spp. MDR | Klebsiella spp. R to carbapenems | Klebsiella spp. R to ES cephalosporins | Carbapenem-R Enterobacteriaceae | E. faecalis R to vancomycin | E. faecium R to vancomycin | Enterobacter spp. R to ES cephalosporins |
|---|---|---|---|---|---|---|---|
| E. coli R to fluoroquinolones | 0.44 (0.12,0.68) | 0.41 (0.08,0.66) | 0.42 (0.1,0.67) | 0.43 (0.13,0.66) | 0.49 (0.18,0.71) | 0.07 (-0.33,0.44) | 0.22 (-0.17,0.55) |
| E. coli MDR | 0.64 (0.39,0.81) | 0.71 (0.49,0.85) | 0.67 (0.43,0.82) | 0.74 (0.55,0.86) | 0.57 (0.29,0.77) | 0.27 (-0.13,0.6) | 0.29 (-0.09,0.6) |
| E. coli R to ES cephalosporins | 0.64 (0.38,0.8) | 0.62 (0.35,0.79) | 0.72 (0.51,0.85) | 0.7 (0.49,0.84) | 0.5 (0.18,0.72) | 0.1 (-0.3,0.47) | 0.47 (0.12,0.72) |
| P. aeruginosa R to fluroquinolones | 0.24 (-0.12,0.54) | 0.14 (-0.22,0.46) | 0.28 (-0.07,0.57) | 0.23 (-0.13,0.53) | 0.21 (-0.15,0.52) | -0.08 (-0.46,0.32) | 0.31 (-0.07,0.62) |
| P. aeruginosa R to aminoglycosides | 0.14 (-0.22,0.46) | 0.16 (-0.19,0.47) | 0.19 (-0.16,0.49) | 0.19 (-0.17,0.5) | 0.31 (-0.04,0.59) | -0.05 (-0.43,0.35) | 0.35 (-0.02,0.64) |
| P. aeruginosa MDR | 0.22 (-0.14,0.52) | 0.27 (-0.08,0.55) | 0.22 (-0.13,0.52) | 0.29 (-0.06,0.58) | 0.43 (0.1,0.67) | 0.06 (-0.33,0.44) | 0.28 (-0.11,0.59) |
| P. aeruginosa R to pip/tazobactam | 0.05 (-0.3,0.39) | 0.14 (-0.21,0.46) | 0.05 (-0.29,0.38) | 0.1 (-0.26,0.43) | 0.09 (-0.27,0.42) | 0.11 (-0.29,0.47) | 0.05 (-0.33,0.42) |
| P. aeruginosa R to ES cephalosporins | 0.25 (-0.1,0.54) | 0.28 (-0.06,0.57) | 0.23 (-0.12,0.53) | 0.3 (-0.05,0.58) | 0.43 (0.1,0.67) | 0.25 (-0.15,0.58) | 0.27 (-0.12,0.58) |
| P. aeruginosa (R+I) to carbapenems | 0.18 (-0.17,0.49) | 0.17 (-0.18,0.48) | 0.15 (-0.2,0.47) | 0.2 (-0.15,0.51) | 0.44 (0.11,0.68) | 0.04 (-0.35,0.42) | 0.2 (-0.19,0.53) |

**Table S15**: Correlations between state-specific prevalence of resistance in the CDC Antibiotic Resistance Patient Safety Atlas data [3] between 2011-2014 for certain combinations of antibiotics/bacteria in CAUTI samples in adults aged 19-64y.

| | Klebsiella spp. MDR | Klebsiella spp. R to carbapenems | Klebsiella spp. R to ES cephalosporins | Carbapenem-R Enterobacteriaceae | E. faecalis R to vancomycin | E. faecium R to vancomycin | Enterobacter spp. R to ES cephalosporins |
|---|---|---|---|---|---|---|---|
| Klebsiella spp. MDR | 1 | 0.89 (0.79,0.94) | 0.96 (0.92,0.98) | 0.87 (0.75,0.93) | 0.33 (-0.02,0.61) | 0.23 (-0.18,0.58) | 0.47 (0.11,0.72) |
| Klebsiella spp. R to carbapenems | 0.89 (0.79,0.94) | 1 | 0.86 (0.73,0.93) | 0.95 (0.91,0.98) | 0.39 (0.06,0.65) | 0.28 (-0.12,0.6) | 0.4 (0.03,0.67) |
| Klebsiella spp. R to ES cephalosporins | 0.96 (0.92,0.98) | 0.86 (0.73,0.93) | 1 | 0.85 (0.72,0.92) | 0.31 (-0.04,0.59) | 0.14 (-0.26,0.5) | 0.55 (0.22,0.77) |
| Carbapenem-R Enterobacteriaceae | 0.87 (0.75,0.93) | 0.95 (0.91,0.98) | 0.85 (0.72,0.92) | 1 | 0.47 (0.15,0.7) | 0.21 (-0.2,0.56) | 0.36 (-0.02,0.65) |
| E. faecalis R to vancomycin | 0.33 (-0.02,0.61) | 0.39 (0.06,0.65) | 0.31 (-0.04,0.59) | 0.47 (0.15,0.7) | 1 | 0.27 (-0.13,0.59) | 0.41 (0.04,0.68) |
| E. faecium R to vancomycin | 0.23 (-0.18,0.58) | 0.28 (-0.12,0.6) | 0.14 (-0.26,0.5) | 0.21 (-0.2,0.56) | 0.27 (-0.13,0.59) | 1 | 0.07 (-0.34,0.46) |
| Enterobacter spp. R to ES cephalosporins | 0.47 (0.11,0.72) | 0.4 (0.03,0.67) | 0.55 (0.22,0.77) | 0.36 (-0.02,0.65) | 0.41 (0.04,0.68) | 0.07 (-0.34,0.46) | 1 |

**Table S16**: Correlations between state-specific prevalence of resistance in the CDC Antibiotic Resistance Patient Safety Atlas data [3] between 2011-2014 for certain combinations of antibiotics/bacteria in CAUTI samples in adults aged 19-64y.

**Section S4:** Correlations between prevalence of antibiotic resistance and rates of hospitalization with septicemia

Tables S17-S21 exhibit, for the different age groups of adults, the linear (Pearson) and Spearman correlations between the state-specific prevalence (percentages) of antibiotic resistance for the different combinations of antibiotics/bacteria in the age-specific catheter-associated urinary tract infection (CAUTI) samples in the CDC AR Atlas data [3], 2011-14 and the state-specific average annual rates of hospitalizations between 2011-12 with septicemia in the diagnosis recorded in the HealthCare Cost and Utilization Project (HCUP) data [1] per 100,000 individuals in the corresponding age group. For each age group, results are presented for the combinations of antibiotics/bacteria for which at least 10 states reported the corresponding data. Additionally, Tables S17-S21 exhibit for each age group and combination of bacteria/antibiotics (i) the average state-specific prevalence of resistance in the CAUTI samples, (ii) the number of states reporting the corresponding data.

| Combination of bacteria/antibiotics | Linear Correlation | Spearman rho (p-value) | Average resistance prevalence | States reporting |
|---|---|---|---|---|
| E.coli R to fluoroquinolones | 0.7 (0.5,0.83) | 0.72 (0.0000003) | 29.9% | 42 |
| E.coli MDR | 0.65 (0.43,0.79) | 0.64 (0.000006) | 6.2% | 42 |
| Klebsiella spp. R to ES cephalosporins | 0.54 (0.26,0.74) | 0.54 (0.0007) | 14.3% | 36 |
| E.coli R to ES cephalosporins | 0.54 (0.27,0.73) | 0.47 (0.002) | 11.7% | 40 |
| Klebsiella spp. R to carbapenems | 0.49 (0.19,0.7) | 0.62 (0.00005) | 5.2% | 36 |
| Klebsiella spp. MDR | 0.49 (0.19,0.7) | 0.56 (0.0004) | 9.8% | 36 |
| Enterobacter spp. R to ES cephalosporins | 0.48 (0.16,0.72) | 0.52 (0.003) | 31.8% | 31 |
| Carbapenem-Resistant Enterobacteriaceae | 0.42 (0.12,0.65) | 0.50 (0.001) | 2.4% | 39 |
| E. faecalis R to vancomycin | 0.42 (0.1,0.66) | 0.45 (0.006) | 8% | 35 |
| P. aeruginosa R to fluroquinolones | 0.42 (0.11,0.65) | 0.39 (0.02) | 21.7% | 37 |
| P. aeruginosa R to aminoglycosides | 0.39 (0.08,0.64) | 0.48 (0.003) | 8.7% | 37 |
| E. faecium R to vancomycin | 0.38 (-0.03,0.68) | 0.30 (0.15) | 84.7% | 24 |
| P. aeruginosa MDR | 0.30 (-0.02,0.57) | 0.32 (0.06) | 13% | 37 |
| P. aeruginosa R to pip/tazobactam | 0.28 (-0.06,0.57) | 0.36 (0.03) | 8.5% | 34 |
| Enterobacter spp. MDR | 0.21 (-0.16,0.53) | 0.16 (0.41) | 10.7% | 30 |
| MRSA | 0.21 (-0.25,0.59) | 0.18 (0.42) | 60.5% | 21 |
| E.coli R to carbapenems | 0.18 (-0.14,0.47) | 0.23 (0.16) | 0.70% | 39 |
| P. aeruginosa (R+I) to carbapenems | 0.11 (-0.23,0.42) | 0.03 (0.89) | 18.50% | 36 |
| Coagulase-Negative Staph (R+I) to vancomcyin | 0.10 (-0.38,0.54) | 0.17 (0.5) | 1.60% | 18 |
| E. faecalis R to daptomycin | 0.07 (-0.42,0.54) | 0.0 (0.99) | 0.70% | 17 |
| Enterobacter spp. R to carbapenems | 0.01 (-0.37,0.39) | -0.08 (0.7) | 5.90% | 27 |
| P. aeruginosa R to ES cephalosporins | -0.01 (-0.33,0.31) | 0.06 (0.72) | 9.90% | 37 |
| Coagulase-Negative Staph R to vancomcyin | -0.05 (-0.5,0.43) | 0.05 (0.86) | 0.90% | 18 |

**Table S17:** (i) Correlations between the state-specific resistance prevalence in CAUTI samples for hospitalized adults aged 65+y [3], 2011-14 and the state-specific average annual rates of septicemia hospitalizations (principal or secondary diagnosis) reported in [1] per 100,000 adults aged 85+y, 2011-12; (ii) average state-specific prevalence of resistance in the CAUTI samples; (ii) the number of states reporting the corresponding data.

| Combination of bacteria/antibiotics | Linear Correlation | Spearman rho (p-value) | Average resistance prevalence | States reporting |
|---|---|---|---|---|
| E.coli R to fluoroquinolones | 0.67 (0.45,0.81) | 0.74 (0.0000002) | 29.9% | 42 |
| E.coli MDR | 0.64 (0.41,0.79) | 0.64 (0.000006) | 6.2% | 42 |
| Klebsiella spp. R to ES cephalosporins | 0.52 (0.24,0.73) | 0.55 (0.0005) | 14.3% | 36 |
| E.coli R to ES cephalosporins | 0.50 (0.23,0.7) | 0.46 (0.003) | 11.7% | 40 |
| E. faecalis R to vancomycin | 0.50 (0.19,0.71) | 0.51 (0.002) | 8% | 35 |
| Klebsiella spp. R to carbapenems | 0.48 (0.19,0.7) | 0.65 (0.00002) | 5.2% | 36 |
| P. aeruginosa R to fluroquinolones | 0.48 (0.19,0.7) | 0.46 (0.004) | 21.7% | 37 |
| E. faecium R to vancomycin | 0.47 (0.09,0.74) | 0.39 (0.06) | 84.7% | 24 |
| P. aeruginosa R to aminoglycosides | 0.46 (0.16,0.68) | 0.55 (0.0004) | 8.7% | 37 |
| Klebsiella spp. MDR | 0.45 (0.15,0.68) | 0.56 (0.0004) | 9.8% | 36 |
| Carbapenem-Resistant Enterobacteriaceae | 0.44 (0.15,0.67) | 0.56 (0.0002) | 2.4% | 39 |
| Enterobacter spp. R to ES cephalosporins | 0.43 (0.09,0.68) | 0.47 (0.01) | 31.8% | 31 |
| P. aeruginosa MDR | 0.36 (0.04,0.61) | 0.39 (0.02) | 13% | 37 |
| P. aeruginosa R to pip/tazobactam | 0.34 (0,0.61) | 0.44 (0.009) | 8.5% | 34 |
| MRSA | 0.32 (-0.12,0.66) | 0.23 (0.3) | 60.5% | 21 |
| P. aeruginosa (R+I) to carbapenems | 0.16 (-0.18,0.47) | 0.10 (0.5) | 18.5% | 36 |
| E. coli R to carbapenems | 0.16 (-0.16,0.45) | 0.22 (0.17) | 0.7% | 39 |
| Enterobacter spp. MDR | 0.154 (-0.22,0.49) | 0.22 (0.17) | 10.70% | 30 |
| Coagulase-Negative Staph (R+I) to vancomycin | 0.15 (-0.34,0.58) | 0.15 (0.44) | 1.60% | 18 |

| | | | | |
|---|---|---|---|---|
| E. faecalis R to daptomycin | 0.095 (-0.4,0.55) | 0.16 (0.52) | 0.70% | 17 |
| Enterobacter spp. R to carbapenems | 0.083 (-0.31,0.45) | 0.04 (0.88) | 5.90% | 27 |
| Coagulase-Negative Staph R to vancomcyin | 0.013 (-0.46,0.48) | 0.0 (0.99) | 0.90% | 18 |
| P. aeruginosa R to ES cephalosporins | 0.0 (-0.32,0.32) | 0.11 (0.68) | 9.90% | 37 |

**Table S18:** (i) Correlations between the state-specific resistance prevalence in CAUTI samples for hospitalized adults aged 65+y [3], 2011-14 and the state-specific average annual rates of septicemia hospitalizations (principal or secondary diagnosis) reported in [1] per 100,000 adults aged 75-84y, 2011-12; (ii) average state-specific prevalence of resistance in the CAUTI samples; (ii) the number of states reporting the corresponding data.

| Combination of bacteria/antibiotics | Linear Correlation | Spearman rho (p-value) | Average resistance prevalence | States reporting |
|---|---|---|---|---|
| E.coli R to fluoroquinolones | 0.61 (0.38,0.77) | 0.67 (0.000003) | 29.9% | 42 |
| E. faecalis R to vancomycin | 0.57 (0.29,0.76) | 0.53 (0.001) | 8% | 35 |
| E.coli MDR | 0.56 (0.3,0.74) | 0.54 (0.0002) | 6.2% | 42 |
| P. aeruginosa R to fluroquinolones | 0.47 (0.17,0.69) | 0.48 (0.003) | 21.7% | 37 |
| P. aeruginosa R to aminoglycosides | 0.46 (0.16,0.68) | 0.52 (0.0009) | 8.7% | 37 |
| Klebsiella spp. R to ES cephalosporins | 0.43 (0.11,0.66) | 0.48 (0.003) | 14.3% | 36 |
| E. faecium R to vancomycin | 0.41 (0.01,0.7) | 0.31 (0.14) | 84.7% | 24 |
| MRSA | 0.40 (-0.04,0.71) | 0.35 (0.13) | 60.5% | 21 |
| Klebsiella spp. R to carbapenems | 0.38 (0.05,0.63) | 0.51 (0.001) | 5.2% | 36 |
| Carbapenem-Resistant Enterobacteriaceae | 0.37 (0.06,0.61) | 0.49 (0.001) | 2.4% | 39 |
| P. aeruginosa MDR | 0.36 (0.04,0.61) | 0.39 (0.02) | 13% | 37 |
| E. coli R to ES cephalosporins | 0.36 (0.05,0.6) | 0.33 (0.04) | 11.7% | 40 |
| Klebsiella spp. MDR | 0.34 (0.02,0.6) | 0.44 (0.007) | 9.8% | 36 |
| Enterobacter spp. R to ES cephalosporins | 0.31 (-0.05,0.6) | 0.29 (0.12) | 31.8% | 31 |
| P. aeruginosa R to pip/tazobactam | 0.26 (-0.09,0.55) | 0.4 (0.02) | 8.5% | 34 |

| | | | | |
|---|---|---|---|---|
| P. aeruginosa (R+I) to carbapenems | 0.17 (-0.17,0.47) | 0.14 (0.42) | 18.5% | 36 |
| Coagulase-Negative Staph (R+I) to vancomcyin | 0.10 (-0.38,0.54) | 0.08 (0.75) | 1.60% | 18 |
| Enterobacter spp. R to carbapenems | 0.09 (-0.3,0.46) | 0.02 (0.91) | 5.90% | 27 |
| Coagulase-Negative Staph R to vancomycin | 0.09 (-0.39,0.54) | 0.11 (0.66) | 0.90% | 18 |
| E. faecalis R to daptomycin | 0.08 (-0.41,0.54) | 0.05 (0.84) | 0.70% | 17 |
| E. coli R to carbapenems | 0.08 (-0.24,0.39) | 0.17 (0.31) | 0.70% | 39 |
| Enterobacter spp. MDR | 0.0 (-0.36,0.36) | -0.05 (0.80) | 10.70% | 30 |
| P. aeruginosa R to ES cephalosporins | -0.02 (-0.34,0.31) | 0.09 (0.59) | 9.90% | 37 |

**Table S19:** (i) Correlations between the state-specific resistance prevalence in CAUTI samples for hospitalized adults aged 65+y [3], 2011-14 and the state-specific average annual rates of septicemia hospitalizations (principal or secondary diagnosis) reported in [1] per 100,000 adults aged 65-74y, 2011-12; (ii) average state-specific prevalence of resistance in the CAUTI samples; (ii) the number of states reporting the corresponding data.

| Combination of bacteria/antibiotics | Linear Correlation | Spearman rho (p-value) | Average resistance prevalence | States reporting |
|---|---|---|---|---|
| E. coli R to fluoroquinolones | 0.52 (0.25,0.71) | 0.54 (0.0004) | 29.4% | 41 |
| P. aeruginosa MDR | 0.49 (0.19,0.71) | 0.47 (0.005) | 20.5% | 34 |
| P. aeruginosa R to pip/tazobactam | 0.47 (0.16,0.7) | 0.43 (0.011) | 13.4% | 34 |
| P. aeruginosa R to ES cephalosporins | 0.46 (0.14,0.69) | 0.42 (0.014) | 12.7% | 34 |
| P. aeruginosa (R+I) to carbapenems | 0.42 (0.09,0.66) | 0.42 (0.014) | 26.1% | 34 |
| P. aeruginosa R to aminoglycosides | 0.40 (0.07,0.65) | 0.33 (0.05) | 15.2% | 34 |
| E. coli MDR | 0.26 (-0.05,0.53) | 0.33 (0.03) | 5.9% | 41 |
| E. faecalis R to vancomycin | 0.23 (-0.13,0.53) | 0.23 (0.20) | 7.1% | 33 |
| P. aeruginosa R to fluroquinolones | 0.22 (-0.14,0.52) | 0.16 (0.38) | 30.6% | 33 |
| Coagulase-Negative Staph R to vancomycin | 0.19 (-0.22,0.54) | 0.16 (0.43) | 0.5% | 25 |
| Carbapenem-Resistant Enterobacteriaceae | 0.13 (-0.2,0.44) | 0.24 (0.16) | 2.70% | 37 |

| | | | | |
|---|---|---|---|---|
| E. coli R to ES cephalosporins | 0.10 (-0.22,0.4) | 0.16 (0.32) | 11.20% | 40 |
| Enterobacter spp. MDR | 0.07 (-0.31,0.43) | 0.14 (0.48) | 11.30% | 28 |
| Klebsiella spp. R to carbapenems | 0.07 (-0.28,0.4) | 0.17 (0.35) | 6.50% | 34 |
| Enterobacter spp. R to ES cephalosporins | 0.06 (-0.32,0.42) | 0.05 (0.82) | 31.90% | 28 |
| Enterobacter spp. R to carbapenems | 0.05 (-0.35,0.44) | 0.15 (0.49) | 6.10% | 25 |
| Coagulase-Negative Staph (R+I) to vancomcyin | 0.02 (-0.37,0.41) | 0.04 (0.86) | 0.60% | 25 |
| Klebsiella spp. MDR | 0.01 (-0.33,0.34) | 0.05 (0.77) | 12.60% | 34 |
| Klebsiella spp. R to ES cephalosporins | -0.08 (-0.4,0.27) | -0.05 (0.78) | 18.20% | 34 |
| E.coli R to carbapenems | -0.08 (-0.39,0.25) | -0.07 (0.68) | 0.60% | 38 |
| E. faecium R to vancomycin | -0.10 (-0.47,0.3) | -0.08 (0.69) | 87.30% | 26 |
| E. faecalis R to daptomycin | -0.16 (-0.67,0.46) | -0.07 (0.83) | 1.40% | 12 |
| MRSA | -0.39 (-0.75,0.15) | -0.43 (0.11) | 47.50% | 15 |

**Table S20:** (i) Correlations between the state-specific resistance prevalence in CAUTI samples for hospitalized adults aged 19-64y [3], 2011-14 and the state-specific average annual rates of septicemia hospitalizations (principal or secondary diagnosis) reported in [1] per 100,000 adults aged 50-64y, 2011-12; (ii) average state-specific prevalence of resistance in the CAUTI samples; (ii) the number of states reporting the corresponding data.

| Combination of bacteria/antibiotics | Linear Correlation | Spearman rho (p-value) | Average resistance prevalence | States reporting |
|---|---|---|---|---|
| P. aeruginosa R to pip/tazobactam | 0.49 (0.18,0.71) | 0.47 (0.005) | 13.4% | 34 |
| P. aeruginosa R to ES cephalosporins | 0.41 (0.08,0.65) | 0.39 (0.022) | 12.7% | 34 |
| P. aeruginosa MDR | 0.41 (0.08,0.65) | 0.46 (0.007) | 20.5% | 34 |
| P. aeruginosa R to aminoglycosides | 0.36 (0.03,0.63) | 0.36 (0.04) | 15.2% | 41 |
| E. coli R to fluoroquinolones | 0.34 (0.04,0.59) | 0.37 (0.02) | 29.4% | 34 |
| P. aeruginosa (R+I) to carbapenems | 0.29 (-0.05,0.58) | 0.34 (0.048) | 26.1% | 33 |
| P. aeruginosa R to fluroquinolones | 0.16 (-0.2,0.47) | 0.09 (0.63) | 30.6% | 25 |

| | | | | |
|---|---|---|---|---|
| Enterobacter spp. R to carbapenems | 0.14 (-0.27,0.51) | 0.13 (0.53) | 6.1% | 25 |
| Coagulase-Negative Staph R to vancomycin | 0.14 (-0.27,0.5) | 0.19 (0.36) | 0.5% | 41 |
| E. coli MDR | 0.11 (-0.21,0.4) | 0.21 (0.18) | 5.9% | 34 |
| E. faecalis R to vancomycin | 0.06 (-0.29,0.4) | 0.12 (0.51) | 7.10% | 33 |
| Coagulase-Negative Staph (R+I) to vancomycin | 0.02 (-0.38,0.41) | 0.09 (0.67) | 0.60% | 25 |
| Enterobacter spp. MDR | 0.0 (-0.38,0.37) | 0.05 (0.79) | 11.30% | 28 |
| Carbapenem-Resistant Enterobacteriaceae | -0.05 (-0.37,0.28) | 0.08 (0.63) | 2.70% | 37 |
| E. faecium R to vancomycin | -0.07 (-0.44,0.33) | -0.14 (0.49) | 87.30% | 26 |
| E. coli R to ES cephalosporins | -0.07 (-0.37,0.25) | 0.05 (0.76) | 11.20% | 40 |
| Enterobacter spp. R to ES cephalosporins | -0.11 (-0.47,0.27) | -0.05 (0.82) | 31.90% | 28 |
| Klebsiella spp. R to carbapenems | -0.14 (-0.46,0.21) | -0.04 (0.85) | 6.50% | 34 |
| E. coli R to carbapenems | -0.14 (-0.44,0.19) | -0.09 (0.60) | 0.60% | 38 |
| Klebsiella spp. MDR | -0.22 (-0.52,0.12) | -0.17 (0.33) | 12.60% | 34 |
| Klebsiella spp. R to ES cephalosporins | -0.30 (-0.58,0.05) | -0.29 (0.10) | 18.20% | 34 |
| MRSA | -0.36 (-0.74,0.19) | -0.55 (0.04) | 47.50% | 15 |
| E. faecalis R to daptomycin | -0.40 (-0.79,0.23) | -0.39 (0.21) | 1.40% | 12 |

**Table S21:** (i) Correlations between the state-specific resistance prevalence in CAUTI samples for hospitalized adults aged 19-64y [3], 2011-14 and the state-specific average annual rates of septicemia hospitalizations (principal or secondary diagnosis) reported in [1] per 100,000 adults aged 18-49y, 2011-12; (ii) average state-specific prevalence of resistance in the CAUTI samples; (ii) the number of states reporting the corresponding data.

**Section S5:** Correlations between prevalence of antibiotic resistance and rates of mortality with sepsis

Tables S22-S26 exhibit, for the different age groups of adults, the linear (Pearson) and Spearman correlations between the state-specific prevalence (percentages) of antibiotic resistance for the different combinations of antibiotics/bacteria in the age-specific catheter-associated urinary tract infection (CAUTI) samples in the CDC AR Atlas data [3], 2011-14 and the state-specific average annual rates of sepsis mortality between 2013-14 [2] per 100,000 individuals in the corresponding age group. For each age group, results are presented for the combinations of antibiotics/bacteria for which at least 10 states reported the corresponding data. Additionally, Tables S22-S26 exhibit for each age group and combination of bacteria/antibiotics (i) the average state-specific prevalence of resistance in the CAUTI samples, (ii) the number of states reporting the corresponding data.

| Combination of bacteria/antibiotics | Linear Correlation | Spearman rho (p-value) | Average resistance prevalence | States reporting |
|---|---|---|---|---|
| E. faecium R to daptomycin | 0.77 (0.26,0.94) | 0.423 (0.22) | 6.5% | 10 |
| E.coli R to fluoroquinolones | 0.72 (0.55,0.83) | 0.694 (6.50E-08) | 30% | 51 |
| Enterobacter spp. R to ES cephalosporins | 0.62 (0.37,0.79) | 0.602 (0.0001) | 32.2% | 35 |
| P. aeruginosa R to fluroquinolones | 0.5 (0.25,0.69) | 0.486 (0.0008) | 22% | 45 |
| Coagulase-Negative Staph (R+I) to vancomcyin | 0.49 (0.07,0.76) | 0.523 (0.01) | 1.4% | 21 |
| E. faecium R to vancomycin | 0.43 (0.07,0.69) | 0.277 (0.15) | 84.5% | 28 |
| P. aeruginosa R to aminoglycosides | 0.41 (0.13,0.63) | 0.411 (0.005) | 8.7% | 45 |
| E.coli MDR | 0.41 (0.15,0.61) | 0.408 (0.003) | 6% | 51 |
| Klebsiella spp. R to ES cephalosporins | 0.39 (0.11,0.62) | 0.406 (0.006) | 14.4% | 44 |
| E.coli R to ES cephalosporins | 0.39 (0.12,0.6) | 0.347 (0.01) | 11.5% | 49 |
| Klebsiella spp. MDR | 0.38 (0.09,0.61) | 0.469 (0.001) | 9.9% | 44 |
| Carbapenem-Resistant Enterobacteriaceae | 0.38 (0.1,0.6) | 0.438 (0.002) | 2.5% | 48 |
| P. aeruginosa MDR | 0.34 (0.05,0.58) | 0.323 (0.03) | 13% | 45 |
| Enterobacter spp. MDR | 0.33 (0,0.6) | 0.306 (0.07) | 10.7% | 35 |
| Klebsiella spp. R to carbapenems | 0.31 (0.01,0.56) | 0.432 (0.004) | 5.2% | 42 |
| P. aeruginosa R to pip/tazobactam | 0.25 (-0.06,0.52) | 0.276 (0.08) | 8.5% | 40 |
| MRSA | 0.25 (-0.17,0.59) | 0.151 (0.48) | 61.2% | 24 |
| Coagulase-Negative Staph R to vancomcyin | 0.21 (-0.24,0.59) | 0.267 (0.24) | 0.8% | 21 |
| P. aeruginosa (R+I) to carbapenems | 0.19 (-0.12,0.46) | 0.123 (0.43) | 18.3% | 43 |
| E.coli R to carbapenems | 0.14 (-0.15,0.41) | 0.149 (0.31) | 0.7% | 48 |

| | | | | |
|---|---|---|---|---|
| E. faecalis R to vancomycin | 0.14 (-0.17,0.43) | 0.203 (0.20) | 7.9% | 41 |
| P. aeruginosa R to ES cephalosporins | 0.13 (-0.17,0.41) | 0.149 (0.33) | 9.8% | 45 |
| E. faecalis R to daptomycin | 0.01 (-0.43,0.45) | -0.067 (0.78) | 0.9% | 20 |
| MRSA I to vancomycin | -0.03 (-0.55,0.51) | -0.034 (0.91) | 0.3% | 14 |
| Enterobacter spp. R to carbapenems | -0.07 (-0.41,0.29) | -0.17 (0.36) | 5.8% | 31 |

**Table S22:** (i) Correlations between the state-specific resistance prevalence in CAUTI samples for hospitalized adults aged 65+y [3], 2011-14 and the state-specific average annual rates of mortality with sepsis (underlying or contributing cause of death) reported in [2] per 100,000 adults aged 85+y, 2013-14; (ii) average state-specific prevalence of resistance in the CAUTI samples; (ii) the number of states reporting the corresponding data.

| Combination of bacteria/antibiotics | Linear Correlation | Spearman rho (p-value) | Average resistance prevalence | States reporting |
|---|---|---|---|---|
| E.coli R to fluoroquinolones | 0.77 (0.63,0.86) | 0.75 (0) | 30% | 51 |
| Enterobacter spp. R to ES cephalosporins | 0.63 (0.38,0.8) | 0.613 (9E-05) | 32.2% | 35 |
| E. faecium R to daptomycin | 0.63 (-0.01,0.9) | 0.399 (0.25) | 6.5% | 10 |
| P. aeruginosa R to fluroquinolones | 0.54 (0.29,0.72) | 0.54 (0.0002) | 22% | 45 |
| Coagulase-Negative Staph (R+I) to vancomcyin | 0.47 (0.05,0.75) | 0.506 (0.02) | 1.4% | 21 |
| E. faecium R to vancomycin | 0.46 (0.11,0.71) | 0.323 (0.09) | 84.5% | 28 |
| E.coli MDR | 0.42 (0.16,0.62) | 0.383 (0.006) | 6% | 51 |
| Klebsiella spp. MDR | 0.4 (0.12,0.63) | 0.456 (0.002) | 9.9% | 44 |
| Klebsiella spp. R to ES cephalosporins | 0.4 (0.12,0.62) | 0.386 (0.01) | 14.4% | 44 |
| P. aeruginosa R to aminoglycosides | 0.4 (0.12,0.62) | 0.406 (0.006) | 8.7% | 45 |
| E.coli R to ES cephalosporins | 0.39 (0.12,0.6) | 0.352 (0.01) | 11.5% | 49 |
| MRSA | 0.38 (-0.02,0.68) | 0.385 (0.06) | 61.2% | 24 |
| Carbapenem-Resistant Enterobacteriaceae | 0.37 (0.09,0.59) | 0.446 (0.001) | 2.5% | 48 |
| P. aeruginosa MDR | 0.34 (0.06,0.58) | 0.352 (0.018) | 13% | 45 |

| | | | | |
|---|---|---|---|---|
| P. aeruginosa R to pip/tazobactam | 0.29 (-0.02,0.55) | 0.349 (0.027) | 8.5% | 40 |
| Enterobacter spp. MDR | 0.29 (-0.04,0.57) | 0.258 (0.13) | 10.7% | 35 |
| Klebsiella spp. R to carbapenems | 0.29 (-0.01,0.55) | 0.402 (0.008) | 5.2% | 42 |
| P. aeruginosa (R+I) to carbapenems | 0.28 (-0.02,0.54) | 0.2 (0.20) | 18.3% | 43 |
| Coagulase-Negative Staph R to vancomcyin | 0.25 (-0.2,0.62) | 0.268 (0.24) | 0.8% | 21 |
| P. aeruginosa R to ES cephalosporins | 0.19 (-0.1,0.46) | 0.201 (0.19) | 9.8% | 45 |
| E. faecalis R to vancomycin | 0.18 (-0.14,0.46) | 0.189 (0.24) | 7.9% | 41 |
| E.coli R to carbapenems | 0.07 (-0.21,0.35) | 0.059 (0.69) | 0.7% | 48 |
| E. faecalis R to daptomycin | 0.03 (-0.42,0.47) | -0.053 (0.82) | 0.9% | 20 |
| MRSA I to vancomycin | -0.02 (-0.54,0.52) | -0.034 (0.91) | 0.3% | 14 |
| Enterobacter spp. R to carbapenems | -0.13 (-0.47,0.23) | -0.246 (0.18) | 30% | 31 |

**Table S23:** (i) Correlations between the state-specific resistance prevalence in CAUTI samples for hospitalized adults aged 65+y [3], 2011-14 and the state-specific average annual rates of mortality with sepsis (underlying or contributing cause of death) reported in [2] per 100,000 adults aged 75-84y, 2013-14; (ii) average state-specific prevalence of resistance in the CAUTI samples; (ii) the number of states reporting the corresponding data.

| Combination of bacteria/antibiotics | Linear Correlation | Spearman rho (p-value) | Average resistance prevalence | States reporting |
|---|---|---|---|---|
| E.coli R to fluoroquinolones | 0.77 (0.63,0.86) | 0.754 (0) | 30% | 51 |
| Enterobacter spp. R to ES cephalosporins | 0.67 (0.43,0.82) | 0.63 (5E-05) | 32.2% | 35 |
| P. aeruginosa R to fluroquinolones | 0.56 (0.32,0.73) | 0.549 (0.0001) | 22% | 45 |
| E. faecium R to vancomycin | 0.53 (0.2,0.76) | 0.43 (0.022) | 84.5% | 28 |
| E. faecium R to daptomycin | 0.53 (-0.16,0.87) | 0.509 (0.13) | 6.5% | 10 |
| MRSA | 0.52 (0.15,0.76) | 0.465 (0.02) | 61.2% | 24 |
| P. aeruginosa R to aminoglycosides | 0.45 (0.18,0.66) | 0.471 (0.001) | 8.7% | 45 |
| Coagulase-Negative Staph (R+I) to vancomycin | 0.41 (-0.03,0.72) | 0.413 (0.06) | 1.4% | 21 |

| | | | | |
|---|---|---|---|---|
| E.coli MDR | 0.4 (0.14,0.61) | 0.408 (0.003) | 6% | 51 |
| P. aeruginosa MDR | 0.38 (0.1,0.61) | 0.389 (0.008) | 13% | 45 |
| Klebsiella spp. R to ES cephalosporins | 0.38 (0.09,0.61) | 0.444 (0.003) | 14.4% | 44 |
| Coagulase-Negative Staph R to vancomcyin | 0.37 (-0.08,0.69) | 0.375 (0.09) | 0.8% | 21 |
| Klebsiella spp. MDR | 0.36 (0.07,0.59) | 0.491 (0.0007) | 9.9% | 44 |
| E.coli R to ES cephalosporins | 0.3 (0.02,0.53) | 0.31 (0.03) | 11.5% | 49 |
| Carbapenem-Resistant Enterobacteriaceae | 0.27 (-0.01,0.52) | 0.425 (0.003) | 2.5% | 48 |
| P. aeruginosa (R+I) to carbapenems | 0.27 (-0.04,0.52) | 0.179 (0.25) | 18.3% | 43 |
| Enterobacter spp. MDR | 0.26 (-0.08,0.55) | 0.233 (0.18) | 10.7% | 35 |
| Klebsiella spp. R to carbapenems | 0.25 (-0.06,0.51) | 0.397 (0.01) | 5.2% | 42 |
| P. aeruginosa R to pip/tazobactam | 0.24 (-0.08,0.51) | 0.312 (0.05) | 8.5% | 40 |
| E. faecalis R to vancomycin | 0.2 (-0.11,0.48) | 0.272 (0.09) | 7.9% | 41 |
| P. aeruginosa R to ES cephalosporins | 0.16 (-0.14,0.43) | 0.16 (0.29) | 9.8% | 45 |
| E. faecalis R to daptomycin | 0.11 (-0.35,0.53) | 0.035 (0.88) | 0.9% | 20 |
| E.coli R to carbapenems | 0.08 (-0.21,0.35) | 0.148 (0.32) | 0.7% | 48 |
| MRSA I to vancomycin | -0.17 (-0.64,0.4) | -0.241 (0.41) | 0.3% | 14 |
| Enterobacter spp. R to carbapenems | -0.2 (-0.51,0.17) | -0.32 (0.08) | 5.8% | 31 |

**Table S24:** (i) Correlations between the state-specific resistance prevalence in CAUTI samples for hospitalized adults aged 65+y [3], 2011-14 and the state-specific average annual rates of mortality with sepsis (underlying or contributing cause of death) reported in [2] per 100,000 adults aged 65-74y, 2013-14; (ii) average state-specific prevalence of resistance in the CAUTI samples; (ii) the number of states reporting the corresponding data.

| Combination of bacteria/antibiotics | Linear Correlation | Spearman rho (p-value) | Average resistance prevalence | States reporting |
|---|---|---|---|---|
| E.coli R to fluoroquinolones | 0.7 (0.52,0.82) | 0.717 (3E-08) | 29.5% | 50 |
| P. aeruginosa MDR | 0.49 (0.21,0.69) | 0.501 (0.0008) | 20.5% | 41 |

| | | | | |
|---|---|---|---|---|
| P. aeruginosa R to ES cephalosporins | 0.48 (0.21,0.69) | 0.44 (0.004) | 12.6% | 41 |
| P. aeruginosa R to fluroquinolones | 0.4 (0.1,0.63) | 0.347 (0.03) | 31% | 39 |
| Enterobacter spp. MDR | 0.37 (0.02,0.63) | 0.404 (0.02) | 11.9% | 32 |
| E.coli MDR | 0.36 (0.09,0.58) | 0.482 (0.0004) | 6% | 50 |
| P. aeruginosa R to pip/tazobactam | 0.36 (0.05,0.61) | 0.376 (0.018) | 13.4% | 39 |
| P. aeruginosa (R+I) to carbapenems | 0.34 (0.03,0.59) | 0.306 (0.06) | 26.1% | 39 |
| P. aeruginosa R to aminoglycosides | 0.33 (0.02,0.58) | 0.306 (0.05) | 14.8% | 41 |
| Enterobacter spp. R to ES cephalosporins | 0.23 (-0.13,0.54) | 0.2 (0.27) | 32.4% | 32 |
| E. faecalis R to daptomycin | 0.23 (-0.32,0.66) | 0.193 (0.49) | 1.4% | 15 |
| Carbapenem-Resistant Enterobacteriaceae | 0.21 (-0.09,0.47) | 0.331 (0.03) | 2.7% | 45 |
| Klebsiella spp. MDR | 0.21 (-0.11,0.48) | 0.267 (0.09) | 12.5% | 42 |
| E. faecalis R to vancomycin | 0.18 (-0.14,0.47) | 0.265 (0.10) | 6.8% | 39 |
| E.coli R to ES cephalosporins | 0.18 (-0.11,0.44) | 0.243 (0.09) | 11.3% | 49 |
| E. faecium R to vancomycin | 0.17 (-0.2,0.5) | 0.189 (0.32) | 87.5% | 30 |
| Klebsiella spp. R to ES cephalosporins | 0.15 (-0.16,0.43) | 0.168 (0.29) | 18% | 42 |
| Klebsiella spp. R to carbapenems | 0.15 (-0.17,0.44) | 0.297 (0.059) | 6.7% | 41 |
| Coagulase-Negative Staph (R+I) to vancomcyin | 0.13 (-0.25,0.47) | 0.021 (0.92) | 0.7% | 29 |
| Coagulase-Negative Staph R to vancomycin | 0.09 (-0.28,0.45) | 0.018 (0.93) | 0.4% | 29 |
| MRSA | 0.09 (-0.4,0.53) | -0.084 (0.74) | 48.2% | 18 |
| Enterobacter spp. R to carbapenems | -0.04 (-0.4,0.33) | -0.04 (0.84) | 6.2% | 29 |
| E.coli R to carbapenems | -0.16 (-0.43,0.14) | -0.052 (0.73) | 0.6% | 46 |

**Table S25:** (i) Correlations between the state-specific resistance prevalence in CAUTI samples for hospitalized adults aged 19-64y [3], 2011-14 and the state-specific average annual rates of mortality with sepsis (underlying or contributing cause of death) reported in [2] per 100,000 adults aged 50-64y, 2013-14; (ii) average state-specific prevalence of resistance in the CAUTI samples; (ii) the number of states reporting the corresponding data.

| Combination of bacteria/antibiotics | Linear Correlation | Spearman rho (p-value) | Average resistance prevalence | States reporting |
|---|---|---|---|---|
| E.coli R to fluoroquinolones | 0.56 (0.34,0.73) | 0.619 (3E-06) | 29.5% | 50 |
| P. aeruginosa MDR | 0.43 (0.14,0.65) | 0.466 (0.002) | 20.5% | 41 |
| P. aeruginosa R to fluroquinolones | 0.4 (0.1,0.63) | 0.34 (0.035) | 12.6% | 41 |
| P. aeruginosa R to ES cephalosporins | 0.38 (0.09,0.62) | 0.345 (0.027) | 31% | 39 |
| Enterobacter spp. MDR | 0.33 (-0.02,0.61) | 0.334 (0.06) | 11.9% | 32 |
| P. aeruginosa R to pip/tazobactam | 0.31 (0,0.57) | 0.307 (0.057) | 6% | 50 |
| P. aeruginosa R to aminoglycosides | 0.31 (0,0.56) | 0.299 (0.058) | 13.4% | 39 |
| P. aeruginosa (R+I) to carbapenems | 0.29 (-0.03,0.55) | 0.238 (0.14) | 26.1% | 39 |
| MRSA | 0.26 (-0.24,0.65) | 0.075 (0.77) | 14.8% | 41 |
| E. faecium R to vancomycin | 0.24 (-0.13,0.55) | 0.171 (0.37) | 32.4% | 32 |
| Enterobacter spp. R to ES cephalosporins | 0.23 (-0.13,0.54) | 0.229 (0.21) | 1.4% | 15 |
| Coagulase-Negative Staph (R+I) to vancomcyin | 0.19 (-0.19,0.52) | 0.099 (0.61) | 2.7% | 45 |
| E.coli MDR | 0.15 (-0.14,0.41) | 0.305 (0.03) | 12.5% | 42 |
| Coagulase-Negative Staph R to vancomcyin | 0.12 (-0.26,0.46) | 0.118 (0.54) | 6.8% | 39 |
| E. faecalis R to vancomycin | 0.07 (-0.25,0.37) | 0.216 (0.19) | 11.3% | 49 |
| Carbapenem-Resistant Enterobacteriaceae | 0.02 (-0.27,0.31) | 0.227 (0.13) | 87.5% | 30 |
| Klebsiella spp. MDR | 0.01 (-0.3,0.31) | 0.177 (0.26) | 18% | 42 |
| Klebsiella spp. R to carbapenems | -0.03 (-0.33,0.28) | 0.212 (0.18) | 6.7% | 41 |
| E.coli R to ES cephalosporins | -0.03 (-0.31,0.25) | 0.07 (0.63) | 0.7% | 29 |
| E. faecalis R to daptomycin | -0.04 (-0.54,0.49) | 0.02 (0.94) | 0.4% | 29 |
| Klebsiella spp. R to ES cephalosporins | -0.05 (-0.35,0.26) | 0.067 (0.67) | 48.2% | 18 |
| Enterobacter spp. R to carbapenems | -0.1 (-0.45,0.28) | -0.083 (0.67) | 6.2% | 29 |

| | | | | |
|---|---|---|---|---|
| E.coli R to carbapenems | -0.19 (-0.45,0.11) | -0.061 (0.69) | 0.6% | 46 |

**Table S26:** (i) Correlations between the state-specific resistance prevalence in CAUTI samples for hospitalized adults aged 19-64y [3], 2011-14 and the state-specific average annual rates of mortality with sepsis (underlying or contributing cause of death) reported in [2] per 100,000 adults aged 18-49y, 2013-14; (ii) average state-specific prevalence of resistance in the CAUTI samples; (ii) the number of states reporting the corresponding data.
